# Single-trial neural dynamics are dominated by richly varied movements

**DOI:** 10.1101/308288

**Authors:** Simon Musall, Matthew T. Kaufman, Ashley L. Juavinett, Steven Gluf, Anne K. Churchland

## Abstract

When experts are immersed in a task, do their brains prioritize task-related activity? Most efforts to understand neural activity during well-learned tasks focus on cognitive computations and specific task-related movements. We wondered whether task-performing animals explore a broader movement landscape, and how this impacts neural activity. We characterized movements using video and other sensors and measured neural activity using widefield and two-photon imaging. Cortex-wide activity was dominated by movements, especially uninstructed movements, reflecting unknown priorities of the animal. Some uninstructed movements were aligned to trial events. Accounting for them revealed that neurons with similar trial-averaged activity often reflected utterly different combinations of cognitive and movement variables. Other movements occurred idiosyncratically, accounting for trial-by-trial fluctuations that are often considered “noise”. This held true for extracellular Neuropixels recordings in cortical and subcortical areas. Our observations argue that animals execute expert decisions while performing richly varied, uninstructed movements that profoundly shape neural activity.

## Introduction

Cognitive functions, such as perception, attention, and decision-making, are often studied in the context of movements. This is because most cognitive processes will naturally lead to action: pondering where to go or with whom to interact both ultimately lead to movement. Laboratory studies of cognition therefore often rely on the careful quantification of instructed movements (e.g., key press, saccade or spout lick), to track outcomes of cognitive computations. However, studying cognition in the context of movements leads to a well-known challenge: untangling neural activity that is specific to cognition from neural activity related to the instructed reporting movements.

To address this problem, researchers often isolate the time of decision from the time of movement using a delay period^1–6^ or prevent the subject from knowing which movement to use until late in the trial^7–9^. A second strategy is to evaluate and account for the extent to which putative cognitive activity is modulated by metrics of the instructed movement: for instance, assessing how decision-related activity co-varies with saccade parameters^2,10,11^ or orienting movements^5^. However, instructed movements are only a subset of possible movements that may modulate neural activity.

In the absence of a behavioral task, uninstructed movements, like running on a wheel, can drive considerable neural activity, even in sensory areas^12–18^. Despite this, uninstructed movements, such as hindlimb flexions during a lick task, are usually not considered when analyzing neural data, exposing two implicit assumptions (Fig. 1A). The first assumption is that uninstructed movements have a negligible impact on neural activity compared to task-related activity or instructed movements. The second assumption is that uninstructed movements occur infrequently and at random times, while instructed movements are task-aligned, occurring at stereotyped moments on each trial. In this case, uninstructed movements would increase neural variability trial-to-trial but their effects could be removed by trial averaging^19^. Both assumptions are largely untested, however, in part due to the difficulty in accurately measuring multiple movement types and systematically relating them to neural activity.

Revealing the impact of uninstructed movements on neural activity is critical for behavioral experiments: movements may account for considerable trial-to-trial variance and may also mimic or overshadow the cognitive processes that are commonly studied in trial-averaged data. It is also unclear whether movements may drive substantial activity in specific areas (such as association or motor areas) but have negligible effects in others. We therefore leveraged multiple movement sensors and dual video recordings to track a wide array of movements and measure their impact on neural activity across the dorsal cortex. A linear encoding model demonstrated that cortex-wide neural variability was dominated by movements, with uninstructed movements outpacing the predictive power of instructed movements and task variables. Both instructed and uninstructed movements also accounted for a large degree of task-aligned activity that was still present after trial-averaging. Our analysis allowed us to separate out such movement-related activity, thereby recovering task-related dynamics that were otherwise obscured. Taken together, these results argue that during cognition, animals engage in a diverse array of uninstructed movements, with profound consequences for neural activity.

## Results

### Cortex-wide imaging during auditory and visual decision-making

We trained mice to report the spatial position of 0.6-s long sequences of auditory click sounds or visual LED stimuli. Animals grabbed handles to initiate trials; stimuli were presented at randomized times after the handle grab (Fig. 1B). Each stimulus was presented twice and after a 1s delay, animals reported a decision and received a water reward for licking the spout corresponding to the stimulus presentation side (Fig. 1C). Animals were trained on either auditory or visual stimuli and achieved expert performance in their trained modality (Fig. 1D). This enabled us to analyze cortical activity in the same detection behavior with different sensory stimuli.

To measure cortex-wide neural dynamics, we used a custom-built widefield macroscope^20^. Mice were transgenic, expressing the Ca^2+^-indicator GCaMP6f in excitatory neurons. Fluorescence was measured through the cleared, intact skull^21^ (Fig. 1E). Using excitation light at two different wavelengths, we isolated Ca^2+^-dependent fluorescence and corrected for intrinsic signals (e.g., hemodynamic responses)^22,23^. The imaging data were then aligned to the Allen Mouse Common Coordinate Framework v3 (CCF, Extended Data Fig. 1). To confirm accurate CCF alignment, we performed retinotopic visual mapping^24,25^ in each animal and found high correspondence between functionally identified visual areas and the CCF (Extended Data Fig. 2).

**Figure 1.**
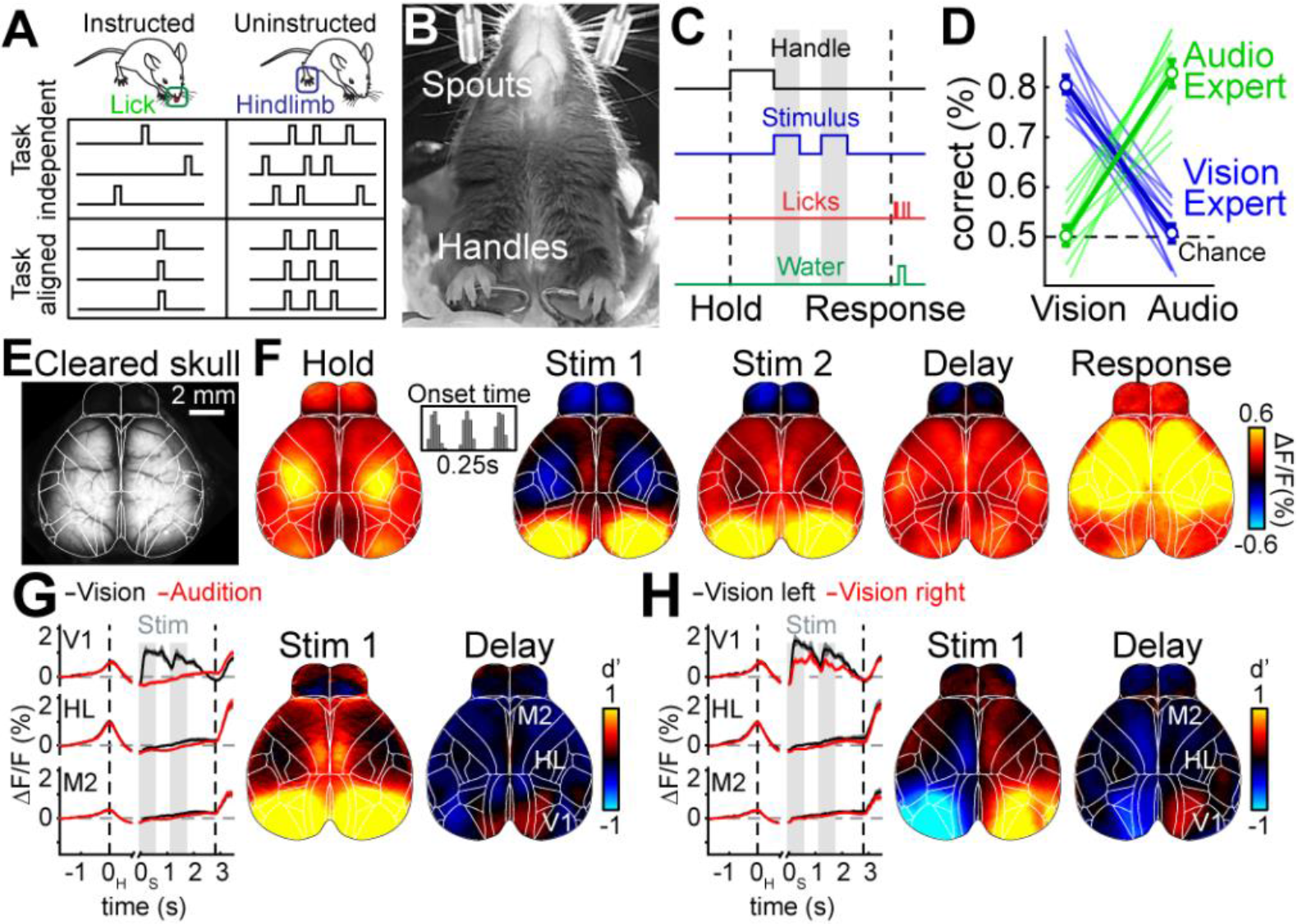
Widefield calcium imaging during auditory and visual decision making. **(A)** Schematic for the two main questions, addressed in this work. First: How important are uninstructed movements compared to instructed movements for driving neural activity during a behavioral task (columns)? Second: To what extent are instructed and uninstructed movement temporally aligned to the task (rows)? Uninstructed movements are exemplified as ‘hindlimb’ but numerous movements are considered throughout this work. **(B)** Bottom view of a mouse in the behavioral setup. **(C)** Single-trial timing of behavior. Mice held the handles for 1s (±0.25s) to trigger the stimulus sequence. One second after stimulus end, water spouts moved towards the mice so they could report a choice. **(D)** Visual experts (blue) had high performance with visual but chance performance with auditory stimuli. Auditory experts (green) showed the opposite. Thin lines show individual animals, thick show averages. Error bars represent mean±s.e.m. **(E)** Example image of cortical surface after skull clearing. Overlaid white lines show Allen atlas borders. **(F)** Cortical activity during different task episodes averaged over 11 mice. Shown are responses when holding the handles (‘Hold’), visual stimulus presentation (‘Stim 1&2)), the subsequent delay (‘Delay’) and the response period (‘Response’). In each trial, stimulus onset was pseudo-randomized within a 0.25-s long time window (inset). **(G)** Left: Traces show average responses in primary visual cortex (V1), hindlimb somatosensory cortex (HL) and secondary motor cortex (M2) on the right hemisphere during visual (black) or auditory (red) stimulation. Trial averages are double-aligned to the time of trial initiation (left dashed line) and stimulus onset (gray bars). Right dashed line indicates response period, fills indicate s.e.m. Right: d’ between visual and auditory trials during first visual stimulus and the subsequent delay period. **(H)** Same as (G) but for correct visual trials on the left versus right side.

Baseline-corrected fluorescence (ΔF/F) revealed significant modulation of neural activity across dorsal cortex (Fig. 1F, Video S1; average response to visual trials, 22 sessions from 11 mice). During trial initiation (handle grab), cortical activity was strongest in sensorimotor areas for hind- and forepaw (‘Hold’). The first visual stimulus caused robust activation of visual areas and weaker responses in medial secondary motor cortex (M2) (‘Stim 1’). Activity in anterior cortex increased during the second stimulus presentation (‘Stim 2’) and the delay (‘Delay’). During the response period, neural activity strongly increased throughout dorsal cortex (‘Response’). A comparison of neural activity across conditions confirmed that neural activity was modulated by whether the stimulus was visual or auditory (Fig. 1G) and presented on the left or the right (Fig. 1H). Differences across both conditions were mainly restricted to primary and secondary visual areas and the posteromedial part of M2.

### Movements dominate cortical activity

To assess the impact of movements, we built a linear encoding model^26,27^ that could take into account a large array of instructed and uninstructed movements as well as various task-variables to predict changes in cortex-wide activity over all trials. We designed model predictors in two ways. The first type of predictor allowed the model to learn an event kernel for different types of behavioral events (Fig. 2A). Some events were related to movements, such as licking or grabbing a handle, and others to task variables like animal choice in a given trial or sensory stimulus onset. For uninstructed movements, we leveraged piezo sensors underneath the animal to track hindpaw movements and video data to detect movements of the nose or whisker pad (Fig. 2B). The resulting traces were then thresholded to isolate individual movement events (Fig. 2C). For all events, we used ridge regression to learn a time-varying event kernel that was time-locked to the behavioral event and predicted related changes in cortex-wide activity. Event kernels perform well when the behavioral event is associated with a similar neural response on every repetition and can recapitulate delayed or multiphasic responses (Fig. 2D). Similar approaches are often used for motion-correction in neuroimaging^28^ and have been successful in fitting complex neural responses to sensory or motor events during cognitive tasks^26,27^. Our second type of predictor was formed from analog (continuously-valued) variables. Analog predictors perform well when neural responses scale linearly with the measured variable, but cannot account for delayed or multiphasic responses (Fig. 2E). Our analog predictors included a measure of pupil diameter (Fig. 2B) but also video data from two cameras observing the animal’s face and body. For video data, we applied singular value decomposition (SVD) to extract the 200 highest-variance video dimensions and used them as analog predictors to provide additional information on movements that we had not previously considered^18,29^.

**Figure 2.**
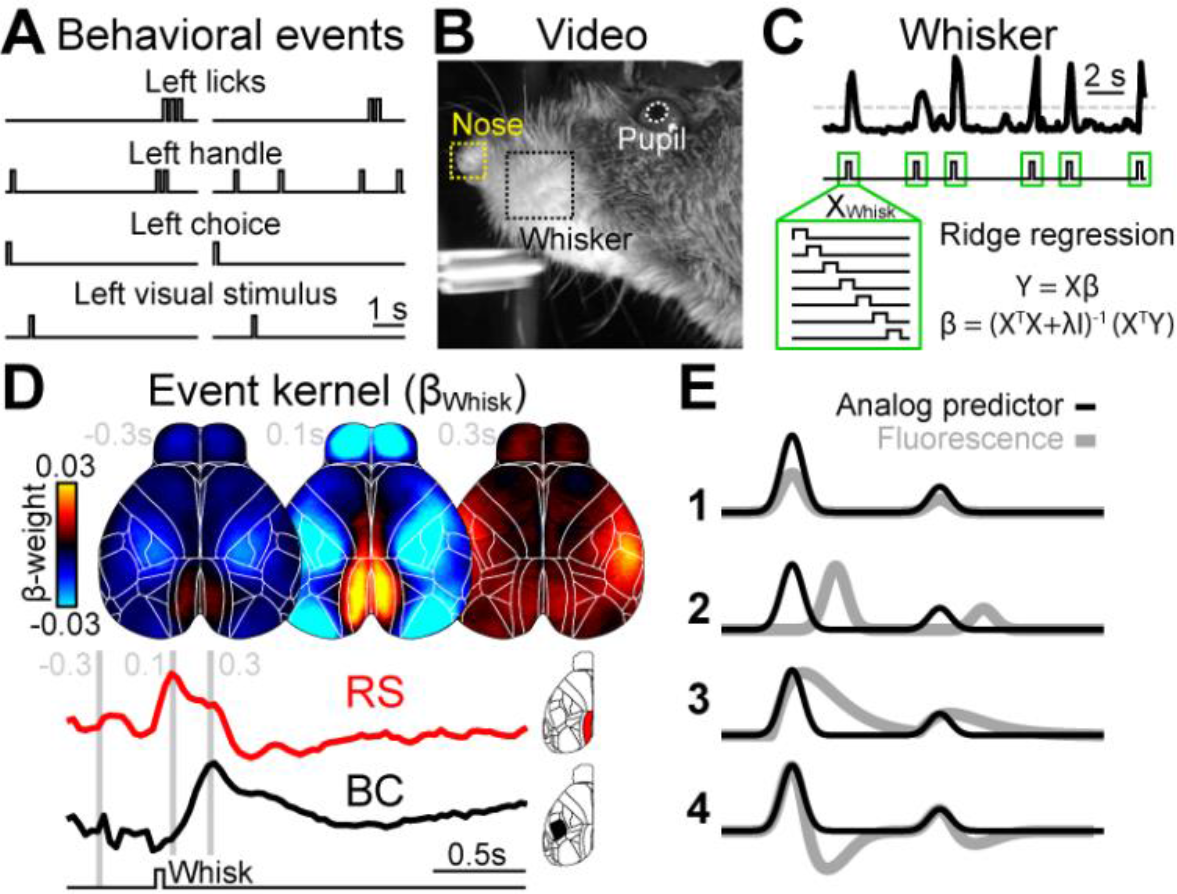
A linear model to reveal behavioral correlates of cortical activity. **(A)** Two example trials, illustrating different classes of behavioral events. **(B)** Image of facial video data with 3 movement variables used in the model. **(C)** Absolute averaged motion energy in the whisker-pad over two trials, showing individual bouts of movement (top trace). Data was thresholded at two standard deviations (dashed line) to infer whisking events. For each event, we created a time-shifted design matrix (X_Whisk_) that was used to compute an event kernel (β_Whisk_) using ridge regression. **(D)** Average β_Whisk_ maps −0.3, 0.1 and 0.3 seconds relative to whisk onset. Whisking caused different responses across cortex, with retrosplenial (RS) being most active 0.1 seconds after whisk onset (bottom red trace) and barrel cortex (BC) after 0.3 seconds (bottom black trace). **(E)** Illustration of an analog predictor (black) fitted to cortical activity (gray). Analog predictors are well-suited to account for correlated changes in activity that scale linearly with a behavioral measure instead of assuming a fixed event response structure (1). In contrast to the event kernel traces in (D), analog predictors cannot account for neural responses that are shifted in time (2) or include additional response features (3-4).

In such a model, it is crucial that the predictors are sufficiently decorrelated. This was accomplished through several sources of trial-to-trial variability, including an experimenter-controlled randomized delay between the animal grabbing the handles and stimulus onset, as well as natural variability in response times and the number of actions (licking, etc.) performed. To ensure that video regressors were not redundant with other movement regressors, we removed overlapping information from the video by orthogonalizing the video regressors to other movement variables (see Methods). We also orthogonalized all uninstructed movement variables against instructed movement variables to ensure that uninstructed movement variables only contained information that could not be captured by instructed movements. To reduce computational cost, we used SVD on the imaging data and ridge regression to fit the model to data components.

To evaluate how well the model captured neural activity at different cortical locations, we computed the 10-fold cross-validated R^2^ (cvR^2^) for the full model during different epochs during the trial (Fig. 3A). While some areas were particularly well predicted in specific trial epochs (e.g., V1 during stimulus presentation), there was high predictive power throughout the cortex during all epochs of the trial. For all data (‘Whole trial’), the model predicted 41.2±0.9% (mean±s.e.m., *n*=22 sessions) of all variance across cortex.

**Figure 3.**
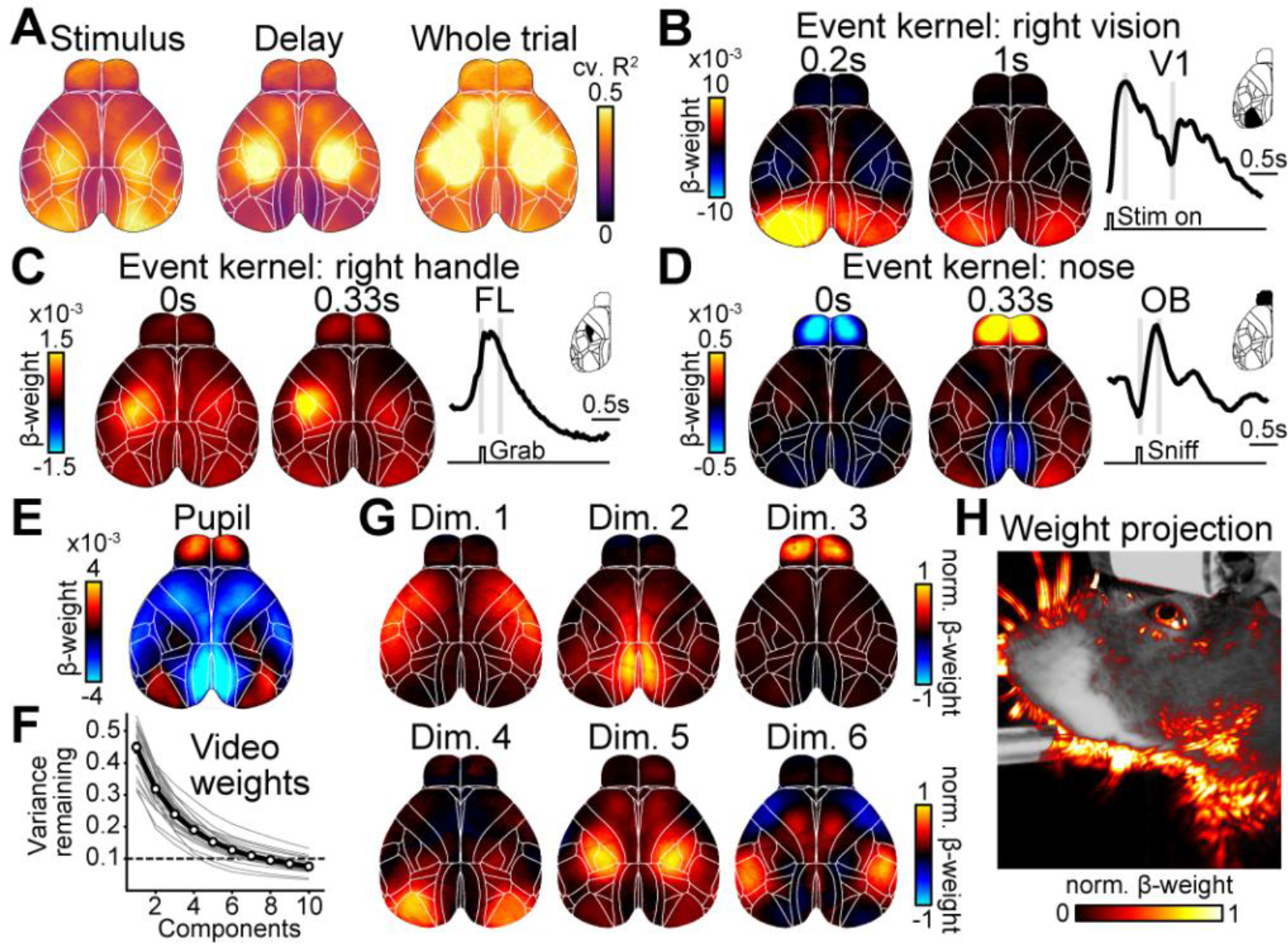
Linear model predicts cortical activity with highly specific model variables. **(A)** Maps of cross-validated explained variance for two task epochs and for the whole trial. **(B)** Example weight maps of the event kernel for right visual stimuli, 0.2 and 1 seconds after stimulus onset. Trace on the right shows, for left V1, the average weights learned by the model. Gray lines indicate times of the maps on the left. **(C)** Same as in (B) but for the event kernel corresponding to right handle grabs, 0 and 0.33 seconds after event onset. **(D)** Same as in (C) but for the event kernel corresponding to nose movements. (**E**) Weight map for the analog pupil regressor. **(F)** Cumulative remaining variance for PCs of model’s weight matrix for all video regressors. Black line shows session average, gray line individual sessions. On average, >90% of all variance was explained by 8 dimensions (dashed line). **(G)** Widefield maps corresponding to the sparsened top 6 video-weight dimensions for an example session. **(H)** Influence of each behavioral video pixel on widefield data. The opacity and color of the overlay were scaled between the 0th and 99th percentile over all values.

Spatiotemporal dynamics of the event kernels sensibly matched known roles of sensory and motor cortices. For example, pixels in left primary visual cortex (V1) were highly positive in response to a rightward visual stimulus (Fig. 3B) with temporal dynamics that matched stimulus responses when averaging over visual trials (Fig. 1H). Left somatosensory (FL) and primary motor forelimb areas were activated during right handle grabs (Fig. 3C) and bi-phasic responses in the olfactory bulb (OB) were associated with nose movements (Fig. 3D). In contrast, the analog pupil regressor was associated with changes in multiple cortical areas, resembling previously observed effects of arousal^30^ (Fig. 3E). To assess the contribution of video data to the model, we performed principal component analysis (PCA) on the model’s weight matrix for all video regressors (Fig. 3F). In many sessions, video weights were at least 8-dimensional, indicating that the video contained different movement patterns that accounted for separate neural responses. To reveal the diversity of relationships between neural responses and video, the first six dimensions of the cortical response weights are shown for an example session, after sparsening (Fig. 3G). When projecting model weights onto video pixels (Fig. 3H), we found that several specific areas of the animal’s face were particularly important, especially around the animal's nose, eye, and jaw. This indicates that video dimensions contributed additional information to the model that was not already contained in other movement variables (e.g., nose movements).

We next sought to address which variables were most important for the model’s success. The simplest way to gauge importance is to fit a model consisting of a single variable and ask how well it predicts the data compared to the full model (Fig. 4A, light green versus both circles combined). However, the cvR^2^ maps for different single-variable models (Fig. 4B, top row) were far less spatially precise than was indicated by their respective event kernels in the full model (Fig. 3, B-D). This is due to overlap in the explanatory power of any given variable with other model variables that contain related information (Fig. 4A, light green inside the black circle). For instance, right visual stimulation also predicted neural activity in the somatosensory mouth area, as well as different parts of anterior cortex. This is because visual stimulation tends to be followed by animal movements and licking in the response period that causes increased activity in corresponding areas. Correspondingly, cvR^2^ maps for the right handle and nose were almost identical, indicating that both might both be associated with other animal movements (Fig. 4B, top right). Single-variable models are therefore an upper bound for the linear information within a given variable. However, the same variable may contribute little to the full model’s overall predictive power if it is largely redundant with other model variables. To measure each variable’s unique contribution to the model, we created reduced models in which we shuffled the regressor set corresponding to a particular variable. The resulting loss of predictive power (ΔR^2^) relative to the full model provides a lower bound for the amount of unique information (non-redundant with any other model variable) that a given variable contributed to the model (Fig. 4A, dark green). In contrast to cvR^2^, ΔR^2^ maps (Fig. 4B, bottom row) were highly spatially localized and matched brain areas that were also most modulated in their corresponding event kernels (Fig. 3). This was consistent in both visual and auditory experts (Extended Data Fig. 3). Control recordings, from animals expressing GFP instead of a GCaMP indicator, confirmed that this was not explained by hemodynamic signals or potential motion artifacts (Extended Data Fig. 4).

We then compared cvR^2^ and ΔR^2^ values (averaged over all pixels) for all model variables. Many variables individually predicted a large amount of neural variance (Fig. 4C, light green bars). Surprisingly, movement variables, both instructed and uninstructed (Fig. 4C, right), contained particularly high predictive power compared to task variables (Fig. 4C, left). Video (‘Video’) and video motion energy (‘Video ME’) were the most predictive model variables, each explaining ~23% of all variance. Differences were even more striking for ΔR^2^ values, which were particularly low for task variables (Fig. 4C, dark green bars). Consider the ‘time’ variable, a regressor set designed to capture signal deviations occurring at consistent times in each trial (similar to an average over all trials). Although the time-only model captured considerable variance (light green bar), eliminating this variable had a negligible effect on the model’s predictive power (dark green bar ~0). This is because other task variables, such as choice or stimulus, could capture time-varying modulation equally well. In contrast, movement variables made much larger unique contributions.

To compare the importance of all instructed or uninstructed movements (Fig. 1A, columns) relative to task variables, we repeated the analysis above on groups of variables (Fig. 4D). Both task and instructed movement groups contained similar predictive power (cvR^2^_Task_=17.6±0.6% cvR^2^_Instructed_=17.3±0.6%) and unique contributions (ΔR^2^_Task_=2.9±0.2% ΔR^2^_Instructed_=3.6±0.2%). Uninstructed movements, however, were more than twice as predictive (cvR^2^_Uninstructed_=38.3±0.9%), with a 5-fold higher unique contribution (ΔR^2^_Uninstructed_=17.8±0.6%). To assess the spatial extent of this large difference, we created pixel-wise ΔR^2^ maps for every group (Fig. 4E). Throughout cortex, unique contributions of uninstructed movements were much higher than either instructed movements or the task group. The contribution of uninstructed movements was particularly prominent in anterior somatosensory and motor cortices, but also clearly evident in posterior areas such as retrosplenial cortex and even V1.

These results strongly suggest that uninstructed movements are critical for predicting neural activity across the dorsal cortex. We considered three possible confounds. First, perhaps the task was insufficiently complex to properly engage cortex. To address this, we tested animals in a more challenging auditory rate discrimination task that is known to require parietal and frontal cortices^31,32^. Results were very similar (Extended Data Fig. 5), arguing that our results were not due to insufficient task complexity. Second, we asked whether the greater predictive power of uninstructed movements might stem from using analog predictors. Indeed, the high ΔR^2^ for video variables shows that analog predictors are particularly effective in detecting movements in an unsupervised manner. To address this, we built a version of the model excluding all analog regressors. Again, uninstructed movements were the most predictive group (Extended Data Fig. 6), demonstrating that their importance does not hinge on the different kinds of model predictors.

Third, we asked whether uninstructed movements might be important because they simply provide additional information about instructed movements (e.g., a preparatory body movement before licking) instead of representing self-generated actions that are independent from the temporal structure of the task. To address this, we computed ΔR^2^ for each group at every time point in the trial (Fig. 4F, Video S2). Here, ΔR^2^_Uninstructed_ was highest in the baseline period when ΔR^2^_Instructed_ was low (black and blue traces). Conversely, ΔR^2^_Uninstructed_ was reduced at times when ΔR^2^_Instructed_ was high, arguing against a tight relation between instructed and uninstructed movements. Instead, their continuously high unique contribution indicates that uninstructed movements mostly represent spontaneous “fidgets” that are independent from instructed movements and the temporal structure of the task.

**Figure 4.**
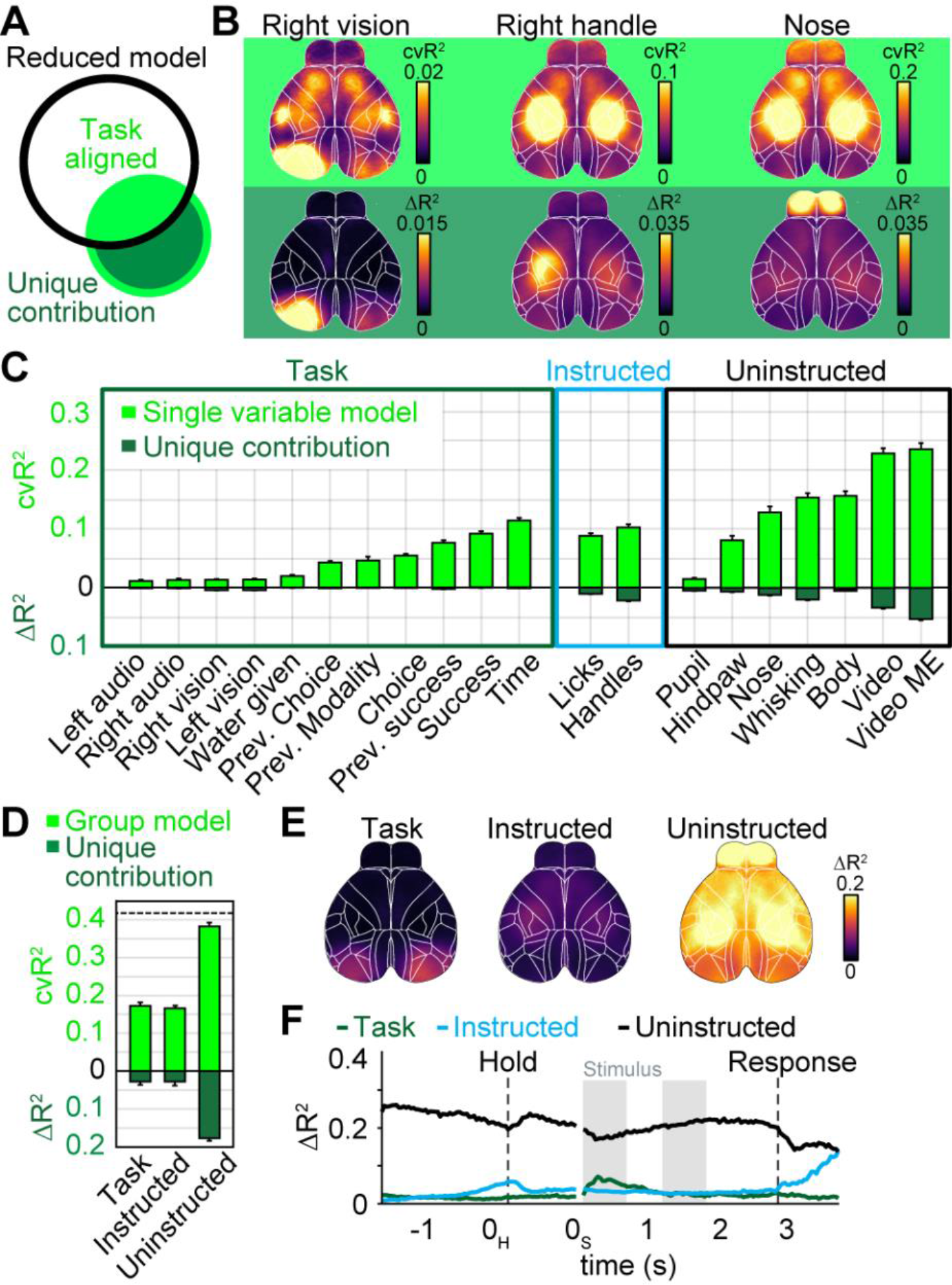
Uninstructed movements dominate cortical activity. **(A)** Black circle denotes information of a reduced model, lacking one variable. The single variable (green circle) has information that overlaps with the reduced model (light green) and a unique contribution (dark green) that increases the full model’s information. **(B)** Top row: Cross-validated explained variance (cvR^2^) maps for different single-variable models. Bottom row: Unique model contribution (ΔR^2^) maps for the same variables. **(C)** Explained variance for single model variables, averaged across cortical maps. Shown is either cvR^2^ (light green) or ΔR^2^ (dark green). Bars represent mean ± s.e.m over 22 sessions. Prev.: previous. **(D)** Explained variance for variable groups. Conventions as in (C). **(E)** ΔR^2^ map for each variable group. **(F)**ΔR^2^ for variable groups at each time-point, averaged across cortex.

### The impact of movements on single-trial and trial-averaged neural activity

To determine how strongly uninstructed movements impacted trial-averaged activity, we asked whether movements were aligned to specific times in the task or only occurred at random times (Fig. 1A, rows). We required instructed movements to occur at specific times (e.g., licking during the response period) and therefore expected them to be strongly task-aligned. Conversely, uninstructed movements like hindpaw flexion might occur at different time points trial-to-trial and therefore be mostly task-independent. If movement timing strongly varies across trials, the hindpaw variable would account for trial-to-trial variability but have no impact on a peri-event time histogram (PETH) that reflects the average over many trials.

To determine the task-aligned and task-independent contributions for each movement variable, we computed the increase in explained variance of the task-only model when a given movement variable was added (Fig. 5A). This is similar to computing unique contributions as described above but here indicates the task-independent contribution (dark blue portion) of a given movement variable. Subtracting this task-independent component from the overall explained variance of each single-movement-variable model (complete blue circle in Fig. 5A, light green bars in Fig. 4C) yields the task-aligned contribution (light blue fill). As expected, instructed movements had a considerable task-aligned contribution, especially the lick variable (Fig. 5B, light blue bars, left). Surprisingly, the same was true for most uninstructed movement variables (Fig. 5B, right), which made considerable task-aligned and task-independent contributions. This was even clearer after pooling variables into groups (Fig. 5C). Here, uninstructed movements contained more task-aligned (cvR^2^_Instructed_=11.0±0.5% cvR^2^_Uninstructed_=14.5±0.5%) and task-independent (ΔR^2^_Instructed_= 6.3±0.3% ΔR^2^_Uninstructed_=23.8±0.7%) explained variance than instructed movements. For both movement groups, the amount of task-aligned information was substantial: instructed and uninstructed movements captured ~63% and ~83% of all information in the task model, respectively.

These results have important implications for interpreting both trial-averaged and single-trial data (e.g., in area M2, Fig. 5D). First, although a task-only model could, by construction, accurately predict the PETH (Fig. 5D, left, top), it cannot account for most single-trial fluctuations (left, bottom 3 rows). Instructed movements also predicted parts of the PETH, in particular at trial times when instructed movements were required (near the black dashed lines, middle top). As with the task model, the instructed movement model failed to predict most single-trial fluctuations (middle, bottom 3 rows). In contrast, an uninstructed movement model also predicted the PETH with good accuracy (right top) and was the only one to also accurately predict neural activity on single trials (right, bottom 3 rows). This demonstrates that much of the single-trial activity that is often assumed to be noise or reflect random network fluctuations is instead related to uninstructed movements.

**Figure 5.**
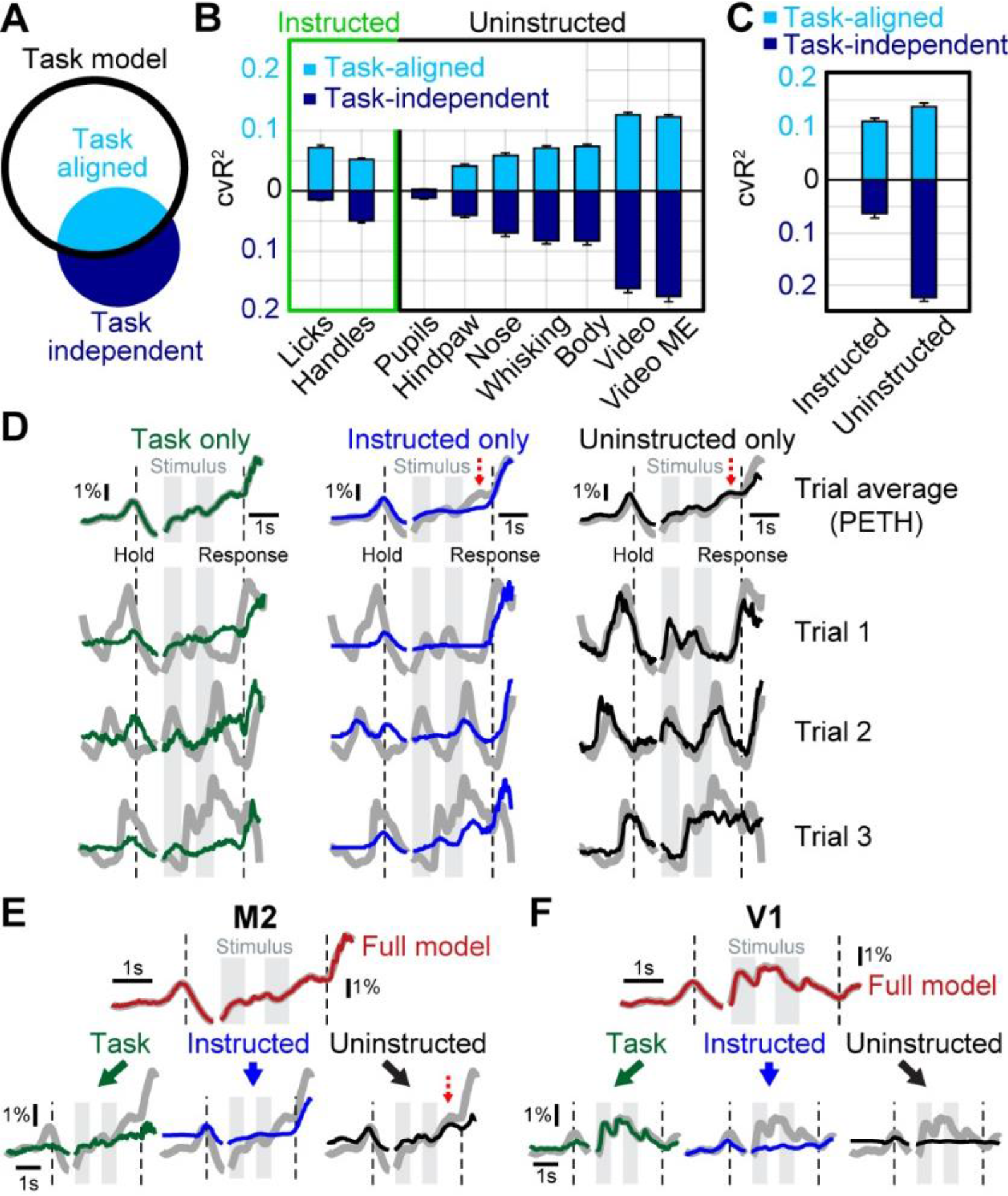
Uninstructed movements make considerable task-aligned and task-independent contributions. **(A)** Black circle denotes task-model information. A movement variable (blue circle) has information that overlaps with the task (task-aligned, light blue) and a unique contribution (dark blue) that is task-independent. **(B)** Explained variance for all movement variables. Shown is task-independent (dark blue) and task-aligned explained variance (light blue). Values averaged across cortex, bars represent mean ± s.e.m over 22 sessions. **(C)** Task-aligned and task-independent explained variance for movement groups. Conventions as in (B). **(D)** Example M2 data from a single animal (gray traces), predicted by different models that were based on a single variable group. PETHs (top row) are mostly well-explained by all three models. Single trials were only well predicted by uninstructed movements (right column, black). **(E)** The M2 PETH is accurately predicted by the full model (top trace, red). Reconstructing the data based on model weights of each variable group allows partitioning the PETH and revealing the groups’ respective contributions to the full model (bottom row). Red dashed arrow indicates an example PETH feature that is best-explained by uninstructed movements. **(F)** Same as in (E) but for V1 data.

Importantly, because many movements are task-aligned, some seemingly cognitive features could be equally well explained by uninstructed movements (Fig. 5D, top row, red dashed arrows). To resolve this issue, we developed an approach to “partition” the PETH into respective contributions from the task, instructed and uninstructed movements, based on how much variance each group accounted for (Fig. 5E). To do so, we fitted the full model to the trial-by-trial data (gray trace). We then split the full model prediction (red trace) into three components based on the weights for each group, without re-fitting. This provides the best estimate of the relative contributions of the task (green traces), instructed movements (blue traces) and uninstructed movements (black traces) to the PETH while explaining the most trial-by-trial neural variance. In M2, partitioning the PETH revealed that diverse dynamics, such as increased activity during the delay (bottom right, red dashed arrow), are best accounted for by uninstructed movements. In contrast, partitioning a PETH in V1 revealed that most features were reassuringly well-explained by the task as they are most likely due to a neural response to visual stimulation (Fig. 5F). This analysis therefore allows a better interpretation of trial-averaged PETHs by using trial-by-trial data to separate task-related activity from the impact of task-aligned movements.

Taken together, these results highlight that most of the activity without trial-averaging is explained by uninstructed movements, which is critical to interpret trial-to-trial variability. Furthermore, uninstructed movements can also occur at predictable time points during a learned behavior and significantly impact the shape of a PETH, sometimes masquerading as activity related to instructed movements or cognitive variables. This reveals a conundrum that is difficult to resolve: many PETH features can be explained by either task or movement variables alone (Fig. 5D top row). Our approach offers insight into this issue: partitioning the PETH into multiple components of neural activity that jointly recreate the complete shape (Fig. 5E, F).

### The impact of movements on single neuron activity

Widefield imaging reflects the pooled activity over many neurons, especially those in superficial layers^22^ and is affected by dendritic and axonal projections^33^. Although we observed substantial area-level specificity (Figs. 3,4B), we further extended our approach to single neurons. Using 2-photon imaging, we measured the activity of over 13,000 single neurons in 10 animals expressing either GCaMP6f (7 mice) or GCaMP6s (3 mice) at depths spanning 150-450μm. We recorded in 5 functionally identified areas, covering large parts of the dorsal cortex. In anterior cortex, we targeted M2 and functionally identified the anterior lateral motor cortex (ALM) and medial motor cortex (MM) based on stereotactic coordinates and their averaged neural responses to licking^3^ (Fig. 6A, top). In posterior cortex we targeted V1 (identified by retinotopic mapping), retrosplenial (RS) and primary somatosensory cortex (S1) (Fig. 6A, bottom). In each area, average population activity was modulated as expected for sensory, motor, and association areas (Fig. 6B).

**Figure 6.**
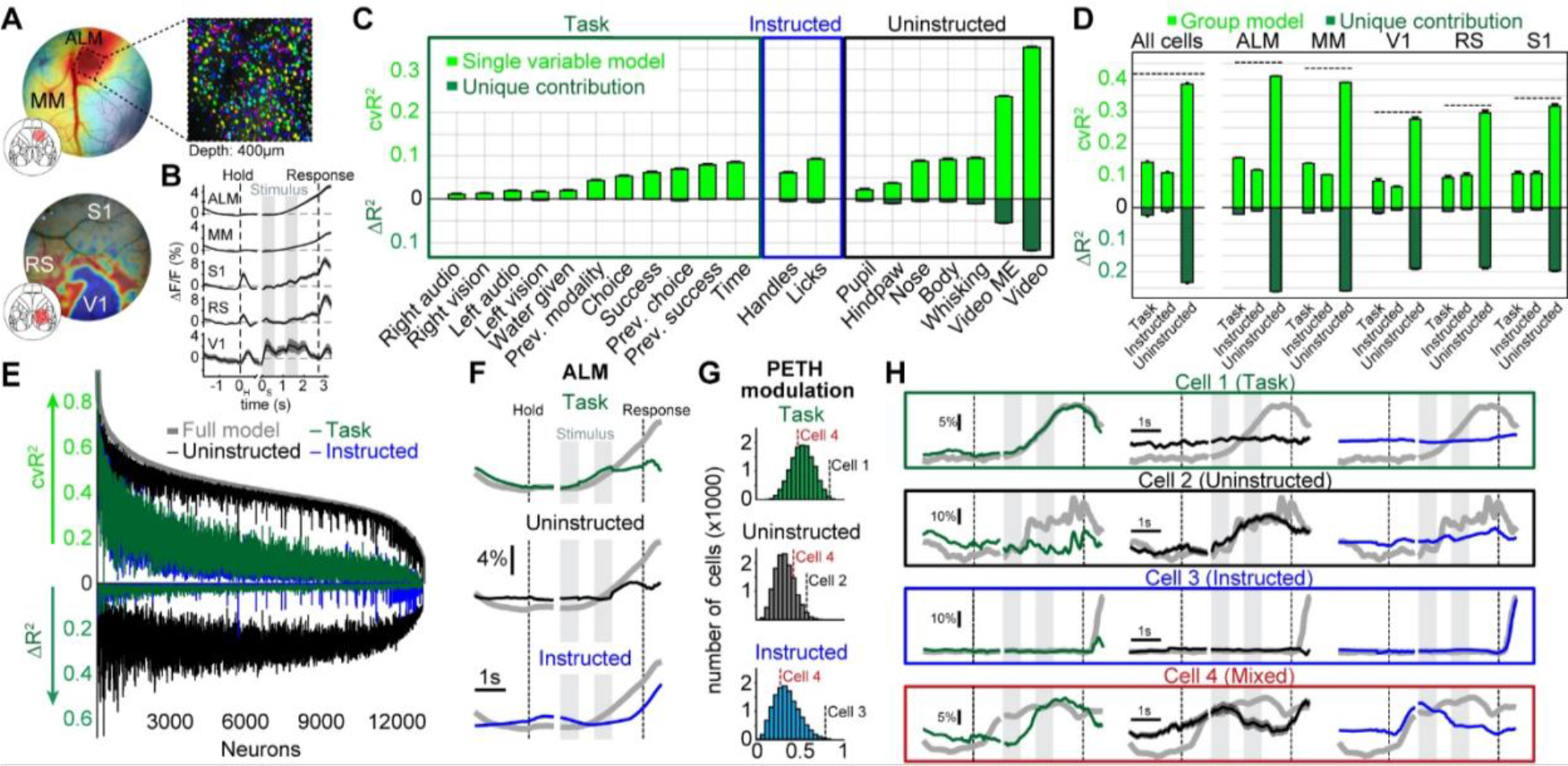
Movements are important for interpreting single-neuron data. **(A)** Example windows in M2 (top) and posterior cortex (bottom). Colors in top window indicate neural response strength during licking. Licking responses match known locations of areas MM and ALM. Colors in bottom window indicate location of visual areas, based on retinotopic mapping. Top right picture shows example field-of-view in ALM with neurons colored randomly. **(B)** Population PETHs from different cortical areas averaged over all trials and recorded neurons. **(C)** Explained variance for single model variables. Shown is either all explained variance (light green) or unique model contribution (dark green). Values averaged over all neurons, bars represent mean ± s.e.m. over 13770 neurons. Prev.: previous. **(D)** Explained variance for variable groups. Conventions as in (C). Left panel shows results for all cells, right panels show results for individual cortical areas. **(E)** Explained variance of variable groups for individual neurons, sorted by full-model performance (light gray trace). Traces above the horizontal axis reflect all explained variance and traces below show unique model contributions (similar to light and dark green bars in D). Colors indicate variable groups. **(F)** Partitioning of the ALM population PETH. Colors show contributions from different variable groups. Summation of all group contributions results in the original PETH (gray traces). **(G)** PETH modulation indices. Histograms show MIs for each variable group. Dashed lines show MI values for example cells in H. **(H)** PETH partitioning of single neurons. Boxes show example cells, most strongly modulated by the task (green), uninstructed movements (black), instructed movements (blue) or a combination of all three (red). Original PETHs in gray.

We then applied our linear model to the 2-photon data, now aiming to predict single cell activity. In the single-cell data, as in the widefield data, the model had high predictive power, capturing 42.8±0.1% (mean±s.e.m., *n*=13770 neurons) of the variance across all neurons. Again, individual movement variables outperformed task variables (Fig. 6C, light green bars) and made larger unique contributions (dark green bars). Given the known causal role of ALM for licking^4,6^, one might expect that the licking variable would be particularly important for predicting single cell activity. Instead, in agreement with our widefield results, almost all movement variables made large contributions. Video-based regressors were again far more powerful than other model variables. The same was true when analyzing cortical areas separately (Extended Data Fig. 7). When grouping variables, we again found that uninstructed movements had much higher predictive power (cvR^2^_Task_=13.9±0.1% cvR^2^_Instructed_=10.1±0.1% cvR^2^_Uninstructed_=38.8±0.1%) and made larger unique contributions (ΔR^2^_Task_=2.8±0.1% ΔR^2^_Instructed_=1.0±0.1% ΔR^2^_Uninstructed_=25.7±0.1%) (Fig. 6D, left). Across areas, full model performance varied from cvR^2^=45.1±0.1% in ALM to cvR^2^=30.0±0.5% in V1 (black dashed lines) potentially due to differences in signal-to-noise ratio across mice. Despite this, the difference between variable groups was highly consistent (Fig. 6D, right). These differences were not driven by outliers but found in almost every recorded neuron (Fig. 6E). Uninstructed movements were consistently the strongest predictors of single neuron activity (Fig. 6E, top black trace) and had the greatest unique contribution to the model (Fig. 6E, bottom black trace). The above results also held true when controlling for potential artifacts due to imaging plane motion by excluding frames with substantial motion (Extended Data Fig. 8).

We again leveraged our ability to partition PETHs into different sources to gain a deeper insight into the functional tuning of single neurons. As with widefield data, partitioning a PETH into different components revealed individual contributions from task, instructed and uninstructed movement variables at different times during the trial. This is exemplified using data averaged over all ALM neurons (Fig. 6F). Average population activity rises consistently over the trial (gray trace), which, at first glance, could be interpreted as a single cognitive process that is progressively increasing. However, the partitioning analysis revealed that this rise reflects the contribution of several components, each with its own dynamics. Task-related activity explained the early rise during the stimulus presentation. It then plateaued during the delay (top green trace). Next, the uninstructed movement contribution increased during the delay (middle black trace) and plateaued before the response period. Finally, the instructed movement contribution increased ~350 ms before the response period. The activity increases contributed by each group are distinct in time and together create average neural activity that smoothly rises over the entire second half of the trial.

We then examined the PETHs of individual cells. To identify cells that were mostly modulated by one variable group versus another, we computed three modulation indices (MIs) for each cell, specifying the extent to which its PETH was best explained by the task, instructed or uninstructed movements (Fig. 6G). Each MI ranges from 0 to 1, with high MI values indicating stronger PETH modulation due to the variable group of interest. Individual cells at the extremes of the MI distributions exemplify strong modulation by task (Fig. 6H, cell 1), uninstructed movements (cell 2) or instructed movements (cell 3). Importantly, the impact of uninstructed movements could not have been inferred from the shape of the PETH alone: while cell 3 was clearly modulated by an instructed movement (licking in the response period), the PETHs of cell 1 and cell 2 exhibited similar temporal dynamics. However, partitioning the PETHs revealed that the activity of these cells reflects very different processes. Notably, this distinction would not have been possible without measuring and analyzing uninstructed movements. PETH partitioning therefore provides additional insight for interpreting the trial average and improves isolation of neurons of interest. Aside from identifying neurons that were purely modulated by a single variable group, we also found that in many cells, we could isolate task-related dynamics that were otherwise overshadowed by the impact of movements on the PETH (cell 4, Extended Data Fig. 9).

Lastly, we tested whether our approach can be extended to other kinds of data by analyzing electrophysiological recordings from Neuropixels probes in visual cortex (VC), superior colliculus (SC) and the midbrain reticular nucleus (MRN) (Fig. 7A). Here, two head-fixed mice were presented with visual looming stimuli and we used PETH partitioning to separate trial-averaged data into stimulus and movement components (Fig. 7B). Consistent with our results from V1 (Fig. 5F), trial-averaged neural responses were well-explained by stimulus predictors with minimal movement contributions. In contrast, across all neurons, trial-by-trial cvR^2^ was far higher for movement variables than for visual stimuli, demonstrating that moment-by-moment neural activity is mostly related to movements (Fig. 7C). This can be seen in an example neuron that exhibited slow fluctuations in firing rate during the recording session (~40 minutes, Fig. 7D), which were not explained by visual stimuli but were highly predictable when using movement regressors. The same was also true when comparing cvR^2^ and ΔR^2^ for stimuli and movements in different cortical and subcortical areas (Fig. 7E). In all recorded brain areas, movements were far more important for predicting neural activity, indicating that the dominance of movements in driving neural activity is not limited to cortex but potentially a brain-wide phenomenon.

**Figure 7.**
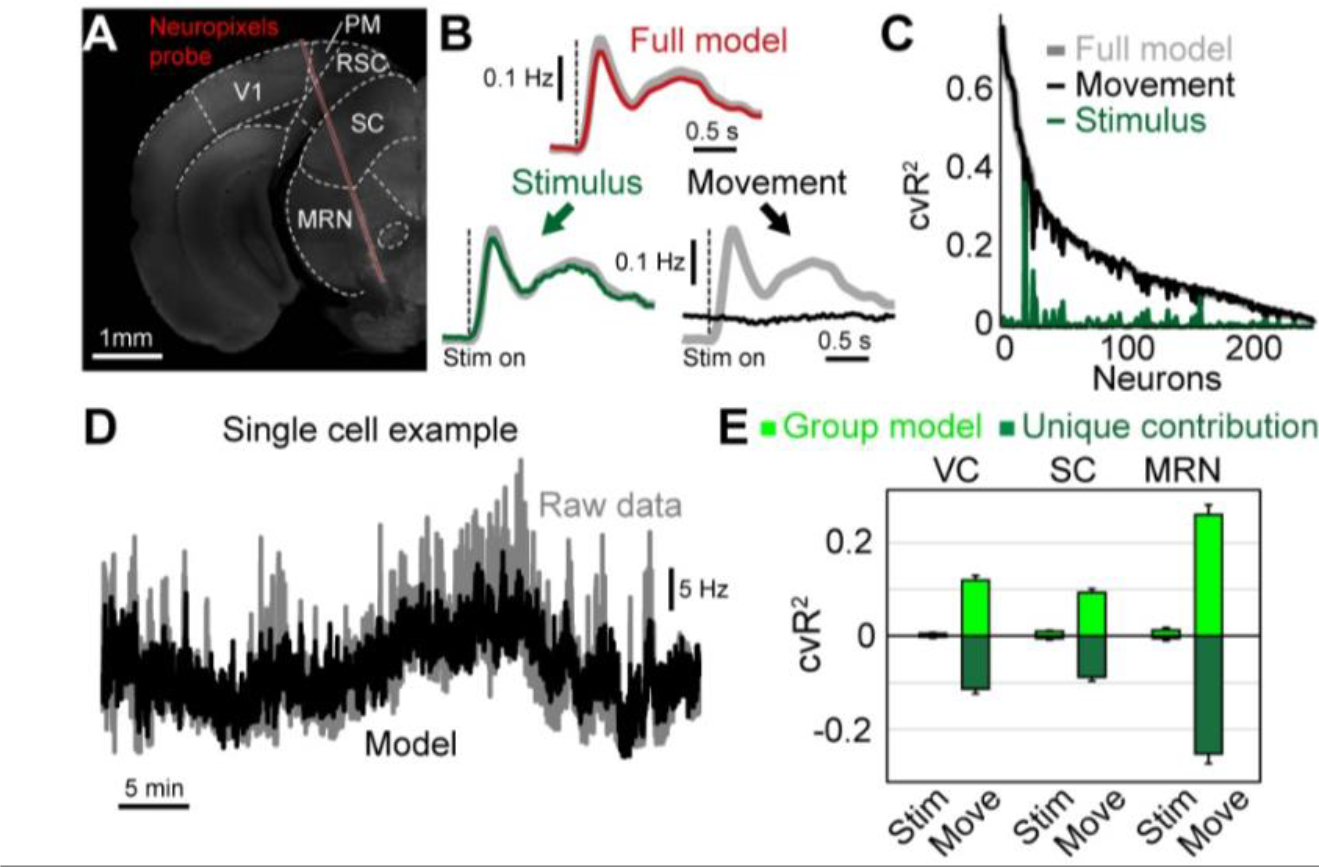
Uninstructed movements predict single-neuron activity in cortical and subcortical areas. **(A)** Coronal slice, −3.8 mm from bregma, showing the location of the Neuropixels probe in an example recording. We recorded single-cell activity from neurons in visual cortex (VC), superior colliculus (SC) and the midbrain reticular nucleus (MRN). **(B)** Population PETH averaged over all neurons and visual stimuli. Gray traces show recorded data. Top red trace shows model reconstruction. Bottom: PETH partitioning of modeled data into stimulus (left, green trace) and movement (right black trace) components. **(C)** Explained variance for all recorded neurons using either the full model (gray) or a movement- (blue) or stimulus-only model (green). **(D)** Activity of a single example neuron over the entire session (~40 minutes). Gray trace is recorded activity, black is the cross-validated model reconstruction. **(E)** Explained variance for stimulus or movement variables in different brain areas. Shown is either all explained variance (light green) or unique model contribution (dark green). Bars represent mean ± s.e.m. over 80, 67 and 85 neurons for VC, SC and MRN, respectively.

## Discussion

Our results demonstrate that cortex-wide neural activity is dominated by movements. By including a wide array of uninstructed movements, our model predicted single-trial neural activity with high accuracy. This was true for thousands of individual neurons, recorded with three different methods, spanning multiple cortical areas and depths. Both instructed and uninstructed movements also had large task-aligned contributions, affecting trial-averaged neural activity. By partitioning PETHs, we separated movement-related from task-related contributions, providing a new tool to unmask truly task-related dynamics and isolate the impact of movements on cortical computations.

Movement-related modulations of neural activity can either reflect the movement itself (due to movement generation, efference copy, or sensory feedback), or changes in internal state that correlate with movements^12,17,34^. Here, we found that movement kernels were highly specific in both cortical areas and temporal dynamics (Fig. 3), indicating that they were probably not strongly reflective of internal state changes. State changes (e.g., arousal) might instead be best captured by analog regressors like the pupil, as indicated by its broad effects across cortex. Video regressors could, in principle, also provide significant internal state information, which may partially explain their particularly high model contributions. However, video weights usually contained at least 8 dimensions that explained neural activity, indicating that it did not solely reflect a low-dimensional state like arousal (Fig. 3). The combined importance of internal states and specific movements highlights the need for tracking multiple movements when assessing their combined impact on neural activity. Using video recordings to track animal movements is a comparably easy way to achieve this goal, especially with new tools that are available for interpreting video data^35–37^, and should therefore become standard practice for physiological recordings.

The vast majority of single neuron activity was well-predicted by uninstructed movements, regardless of brain area or depth (Figs. 6D, 7E, Extended Data Fig. 8). The prevalence of such movement modulation across cortex may explain why apparent task-related activity has been observed in a variety of areas^22,38,39^ and highlights the importance of additional controls, like neural inactivation, to establish involvement in a given behavior^40^. However, in areas with established causal relevance, movements must still be considered. For instance, in ALM, which is known to play a causal role in similar tasks to the one studied here^3,6,21^, many neurons were well-explained by uninstructed movement variables (Fig. 6H). Our partitioning method can help to better interpret such neural activity by identifying neurons that are best explained by task variables. Moreover, for neurons with mixed selectivity for task and movement variables^41^, the model can separate their respective contributions and reveal obscured task-related neural dynamics (Extended Data Fig. 9).

Surprisingly, task variables contained far less predictive power than movement variables, although several areas we studied (V1, A1 and ALM) are known to be involved in similar or simpler detection tasks^21,42^. Importantly, this does not indicate that task-related dynamics are absent from cortical circuits but simply that movements account for a much larger amount of neural variance. Virtually all cortical neurons we recorded also exhibited mixed selectivity and were strongly modulated by movements regardless of whether or not they were also modulated by the task (Fig. 6E). Therefore, even for neurons that are strongly tuned to task variables, cognitive computations are often embedded within a broader context of movement-related information.

The profound impact of uninstructed movements on neural activity suggests that movement signals play a critical role in cortical information processing. Widespread integration of movement signals might be advantageous for sensory processing^15,43^, canceling of self-motion^44^, gating of inputs^45^ or permitting distributed associational learning^46,47^. All of these functions build on the idea that the brain is creating an internal model of the world that is based on the integration of sensory and motor information to predict future events. Our results highlight the notion that such predictive coding is not limited to sensory areas but might be a common theme across cortex^48^: while sensory areas integrate motion signals to predict expected future stimuli^13,49^, motor areas equally integrate sensory signals to detect deviations between intended and actual motor state^50^. The principles of comparing current and predicted feedback also extend to other brain areas like the ventral tegmental area^51^ or the cerebellum^52^. This suggests that robust movement representations may be found throughout the brain.

## Acknowledgements

We thank Onyekachi Odoemene, Sashank Pisupati and Hien Nguyen for technical assistance and scientific discussions, Hongkui Zeng for providing Ai93 mice, Jason Tucciarone and Fred Marbach for breeding assistance, and Alea Mills and Padmina Shrestha for providing GFP mice. Financial support was received from the Swiss National Science foundation (SM, P2ZHP3_161770), the Pew Charitable Trusts (AKC), the Simons Collaboration on the Global Brain (AKC, MTK) and the NIH (EY R01EY022979).

## Methods

### Animal Subjects

The Cold Spring Harbor Laboratory Animal Care and Use Committee approved all animal procedures and experiments. Experiments were conducted with male mice from the ages of 8-25 weeks. All mouse strains were of C57BL/6J background and purchased from Jackson Laboratory. Six transgenic strains were used to create the transgenic mice used for imaging: Emx-Cre (JAX 005628), LSL-tTA (JAX 008600), CaMK2α-tTA (JAX 003010), Ai93 (JAX 024103), G6s2 (JAX 024742) and H2B-eGFP (JAX 006069). Ai93 mice as described below were Ai93; Emx-Cre; LSL-tTA; CaMK2α-tTA. G6s2 as described below were CaMK2α-tTA; tetO GCaMP6s.

For widefield imaging we used 11 Ai93 mice (4 mice were negative for CaMK2α-tTA). Two G6s2 and H2B-eGFP mice were used for widefield control experiments. For two-photon imaging, we recorded from 7 Ai93 mice (6 mice were negative for CaMK2α -tTA) and 3 G6s2 mice. All trained mice were housed in groups of two or more under an inverted 12:12-h light-dark regime and trained during their active dark cycle.

### Surgical procedures

All surgeries were performed under 1-2% isoflurane in oxygen anesthesia. After induction of anesthesia, 1.2 mg/kg of meloxicam was injected subcutaneously and lidocaine ointment was topically applied to the skin. After making a medial incision, the skin was pushed to the side and fixed in position with tissue adhesive (Vetbond, 3M). We then created an outer wall using dental cement (C&B Metabond, Parkell; Ortho-Jet, Lang Dental) while leaving as much of the skull exposed as possible. A circular headbar was attached to the dental cement. For widefield imaging, after carefully cleaning the exposed skull we applied a layer of cyanoacrylate (Zap-A-Gap CA+, Pacer technology) to clear the bone. After the cyanoacrylate was cured, cortical blood vessels were clearly visible.

For 2-photon imaging, instead of clearing the skull, we performed a circular craniotomy using a biopsy punch. For imaging anterior cortex, a 3-mm wide craniotomy was centered 1.5 mm lateral and 1.5 mm anterior to bregma. For imaging posterior cortex, a 4-mm wide craniotomy was centered 1.7 mm lateral and 2 mm posterior to bregma. We then positioned a circular coverslip of similar size over the cortex and sealed the remaining gap between the bone and glass with tissue adhesive. The coverslip window was then secured to the skull using C&B Metabond (Parkell) and the remaining exposed skull was sealed using dental cement. After surgery, animals were kept on a heating mat for recovery and a daily dose of analgesia (1.2 mg/kg Meloxicam) and antibiotics (2.3 mg/kg Enrofloxacin) were administered subcutaneously for at least 3 days or longer.

For electrophysiology experiments, we used 13-17 week old male mice. Mice were given medicated food cups (MediGel CPF, Clear H20 74-05-5022) 1-2 days prior to surgery. We performed a small circular craniotomy over visual cortex using a dental drill. Rather than a circular headbar, a boomerang-shaped custom Titanium headbar was cemented to the skull, just posterior to the eyes, near Bregma. In addition, a small ground screw was drilled into the skull over the cerebellum. The probe was mounted onto a 3D printed piece within an external casing, and affixed to a custom stereotaxic adapter, and lowered into the brain as previously described^53^. Recordings were performed after the mouse had fully recovered from surgery (3-4 days).

### Behavior

The behavioral setup was based on an Arduino-controlled finite state machine (Bpod r0.5, Sanworks) and custom Matlab code (2015b, Mathworks) running on a linux PC. Servo motors (Turnigy TGY-306G-HV), touch sensors and visual stimuli were controlled by microcontrollers (Teensy 3.2, PJRC) running custom code. Twenty-five mice were trained on a delayed two-alternative forced choice (2AFC) spatial discrimination task. Mice initiated trials by touching either of two handles with their forepaws. Handles were mounted on servo motors and were moved out of reach between trials. After one second of holding a handle, sensory stimuli were presented. Sensory stimuli consisted of either a sequence of auditory clicks, or repeated presentation of a visual moving bar (3 repetitions, 200 ms each). Auditory stimuli were presented from either a left or right speaker, and visual stimuli were presented on one of two small LED displays on the left or right side. The sensory stimulus was presented for 600 ms, there was a 500 ms pause with no stimulus, and then the stimulus was repeated for another 600 ms. After the second stimulus, a 1000 ms delay was imposed, then servo motors moved two lick spouts into close proximity of the animal’s mouth. If the animal licked twice to the spout on the same side as the stimulus, he was rewarded with a drop of water. After one spout was contacted twice, the other spout was moved out of reach to force the animal to commit to its decision.

Animals were trained over the course of approximately 30 days. After 2-3 days of restricted water access, animals were head-fixed and received water in the setup. Water was given by presenting a sensory stimulus, subsequently moving the correct spout close to the animal, then dispensing water automatically. After several habituation sessions, animals had to touch the handles to trigger the stimulus presentation. Once animals reliably reached for the handles, the required touch duration was gradually increased up to 1 second. Lastly, the probability for fully self-performed trials, in which both spouts were moved towards the animal after stimulus presentation, was gradually increased until animals reached stable detection performance of 80% or higher.

Each animal was trained exclusively on a single modality (6 visual animals, 5 auditory for widefield imaging; 4 auditory for widefield controls; 5 visual animals, 5 auditory animals for 2-photon imaging). Only during imaging sessions were trials of the untrained modality presented as well. This allowed us to compare neural activity on trials where animals performed stimulus-guided sensorimotor transformations versus trials where animal decisions were random. To ensure that detection performance was not overly affected by presentation of the untrained modality, the trained modality was presented in 75% and the untrained modality in 25% of all trials.

### Behavioral sensors

We used information from several sensors in the behavioral setup to measure different aspects of animal movement. The handles detected contact with the animal’s forepaws, and the lick spouts detected contact with the tongue, using a grounding circuit. An additional piezo sensor (1740, Adafruit LLC) below the animal’s trunk was used to detect hindpaw and whole-body movements. Sensor data were normalized and thresholded at 2 standard deviations to extract hindpaw movements. Based on hindpaw events we created an event-kernel design matrix that was also included in the linear model (see below).

### Video monitoring

Two webcams (C920 and B920, Logitech) were used to monitor animal movements. Cameras were positioned to capture the animal’s face (side view) and the body (bottom view). To target particular behavioral variables of interest, we defined subregions of the video which were then examined in more detail. These included a region surrounding the eye, the whisker pad and the nose. From the eye region we extracted changes in pupil diameter using custom Matlab code. To analyze whisker movements, we computed the absolute temporal derivative averaged over the entire whisker pad. The resulting 1-D trace was then normalized and thresholded at 2 standard deviations to extract whisking events. Based on whisking events we created an event-kernel design matrix that was also included in the linear model (see below). The same approach was used for the nose and pupil diameter.

### Widefield imaging

Widefield imaging was done using an inverted tandem-lens macroscope^20^ in combination with an sCMOS camera (Edge 5.5, PCO) running at 60 fps. The top lens had a focal length of 105 mm (DC-Nikkor, Nikon) and the bottom lens 85 mm (85M-S, Rokinon), resulting in a magnification of 1.24x. The total field of view was 12.5 x 10.5 mm and the image resolution was 640 x 540 pixels after 4x spatial binning (spatial resolution: ~20μm/pixel). To capture GCaMP fluorescence, a 525 nm band-pass filter (#86-963, Edmund optics) was placed in front of the camera. Excitation light was projected on the cortical surface using a 495 nm long-pass dichroic mirror (T495lpxr, Chroma) placed between the two macro lenses. The excitation light was generated by a collimated blue LED (470 nm, M470L3, Thorlabs) and a collimated violet LED (405 nm, M405L3, Thorlabs) that were coupled into the same excitation path using a dichroic mirror (#87-063, Edmund optics). We alternated illumination between the two LEDs from frame to frame, resulting in one set of frames with blue and the other with violet excitation at 30 fps each. Excitation of GCaMP at 405 nm results in non-calcium dependent fluorescence^54^, allowing us to isolate the true calcium-dependent signal by rescaling and subtracting frames with violet illumination from the preceding frames with blue illumination. All subsequent analysis was based on this differential signal at 30 fps.

To ensure that our correction approach effectively removed non-calcium dependent fluorescence, we performed additional control experiments with two GFP-expressing animals (CAG-H2B-eGFP). Here, the hemodynamic correction removed ~90% of variance in the data and the model lost most of its predictive power and spatial specificity (Extended Data Fig. 4). We also imaged two G6s2 mice expressing GCaMP6s^23^. These animals were expected to exhibit stronger calcium-dependent fluorescence due to large signal-to-noise ratio (SNR) of the GCaMP6s indicator. Accordingly, the remaining variance after hemodynamic correction was much higher (Extended Data Fig. 4A-B) and the model predicted even more variance than in Ai93 mice (Extended Data Fig. 4C-D). The β-weights in modulated areas were also strongest with GCaMP6s-expressing mice and close to zero in GFP controls (Extended Data Fig. 4F&H). Together, these controls therefore provide strong evidence that our widefield results were not due to potential contributions from uncorrected hemodynamic signals.

### Two-photon imaging

Two-photon imaging was performed with a resonant-scanning two-photon microscope (Sutter Instruments, Movable Objective Microscope, configured with the “Janelia” option for collection optics), a Ti:Sapphire femtosecond pulsed laser (Ultra II, Coherent Inc.), and a 16X 0.8 NA objective (Nikon Instruments). Images were acquired at 30.9 Hz with an excitation wavelength of 930 nm. All focal planes were between 150-450 μm below the pial surface. The objective height was manually adjusted during the recording in 1-2 μm increments as often as necessary to maintain the same focal plane.

### Preprocessing of neural data

To analyze widefield data, we used SVD to compute the 200 highest-variance dimensions. These dimensions accounted for at least 90% of the total variance in the data. Using 500 dimensions accounted for little additional variance (~0.15%), indicating that additional dimensions were mostly capturing recording noise. SVD returns ‘spatial components’ *U* (of size pixels x components), ‘temporal components’ *V*^T^ (of size components x frames) and singular values *S* (of size components x components) to scale components to match the original data. To reduce computational cost, all subsequent analysis was performed on the product *SV*^T^. *SV*^T^ was high-pass filtered above 0.1Hz using a zero-phase, second-order Butterworth filter. Results of analyses on *SV*^T^ were later multiplied with *U*, to recover results for the original pixel space. All widefield data was rigidly aligned to the Allen Common Coordinate Framework v3, using four anatomical landmarks: the left, center, and right points where anterior cortex meets the olfactory bulbs and the medial point at the base of retrosplenial cortex.

To analyze 2p data we used Suite2P^55^ with model-based background subtraction. The algorithm was used to perform rigid motion correction on the image stack, identify neurons, extract their fluorescence, and correct for neuropil contamination. ΔF/F traces were produced using the method of Jia et al.^56^, skipping the final filtering step. Using these traces, we produced a matrix of size neurons x time, and treated this similarly to *SV*^T^ above. Finally, we confirmed imaging stability by examining the average firing rate of neurons over trials. If all neurons varied substantially at the beginning or end of a session, trials containing the unstable portion were discarded. Recording sessions yielded 188.63±19.28 neurons and contained 378.56±9 performed trials (mean±s.e.m.).

We also used the XY-translation values from the initial motion correction to ask whether motion of the imaging plane (due to imperfect registration or Z-motion) could contribute to our 2-photon results. To account for this possibility, we removed all frames that were translated by more than 2 pixels in the X or Y direction and repeated the analysis (Extended Data Figure 8). In 4983 neurons, we observed negative unique contributions from the task-group, indicative of model overfitting when removing too many data frames. We therefore rejected these neurons from further analysis. For the remaining 8787 neurons, explained variance was highly similar as in our original findings, demonstrating that our results are not explained by motion of the imaging plane.

To compute trial-averages, imaging data were double-aligned to the time when animals initiated a trial and to the stimulus onset. After alignment, single trials consisted of 1.8 s of baseline, 0.83 s of handle touch and 3.3 s following stimulus onset. The randomized additional interval between initiation and stimulus onset (0 - 0.25 s) was discarded in each trial and the resulting trials of equal length were averaged together.

### Neuropixels recordings

To investigate single-neuron responses, a set of linearly expanding (looming, L) or contracting (receding, R) dots (40 cm/s) were presented above the mouse’s head while the mouse was head-restrained but free to move on a wheel^57^. Stimuli were high or low contrast, and either visual only or with an additional auditory looming stimulus -- white noise of increasing volume (80 dB at max volume). Eight different types of stimuli were presented: high and low contrast visual looming (VL), high and low contrast visual receding (VR), high and low contrast audiovisual looming, and high and low contrast audiovisual receding. Stimuli (20 repeats of each type) were 0.5 s long and randomly presented, with randomized gaps (more than 4 s) between stimuli. Results in Fig. 7B were obtained by averaging over all stimulus conditions. A video camera (Basler AG) in combination with infrared lights was used to record the pupil and face during visual stimulation. These videos were synchronized with the electrophysiology data for subsequent analysis.

Electrophysiology data was collected with SpikeGLX (Bill Karsh, https://github.com/billkarsh/SpikeGLX). We recorded from 384 channels spanning ~4 mm of the brain^58^. The data were first median subtracted across channels and time, sorted with Kilosort spike sorting software^59^, and manually curated using phy (https://github.com/kwikteam/phy). Sorted data were analyzed using custom MATLAB code.

### Linear model

The linear model was constructed by combining multiple sets of regressors into a design matrix, to capture signal modulation by different task or motor events (Fig. 2C). Each regressor set (except for ‘analog’ regressors) was structured to capture a time-varying event kernel. To do so, each set included a binary vector containing a pulse at the time of the relevant event, and numerous copies of this vector each shifted in time by one frame relative to the original. For sensory stimuli, we created post-event regressor sets spanning all frames from stimulus onset until the end of the trial. For motor events like licking or whisking, we created peri-event regressor sets that spanned the frames from 0.5 s before until 1 s after each event. Lastly, we created whole-trial regressors, covering each frame in a given trial. Whole-trial regressors were aligned to stimulus onset and contained information about decision variables, such as animal choice or whether a given trial was rewarded. The model also contained several analog regressors, such as 1-D regressors for pupil diameter. To capture animal movements, we used SVD to compute the 200 highest dimensions of video information in both cameras. SVD was performed either on the raw video data (‘video’) or the absolute temporal derivative (motion energy; ‘video ME’). SVD analysis of behavioral video was the same as for the widefield data, and we used the product *SV*^T^ of temporal components and singular values as analog regressors in the linear model. We did not use lagged versions of the analog regressors, including the video regressors.

To use video data regressors, it was important to ensure that their explanatory power did not overlap with other model variables like licking and whisking that can also be inferred from video data. To accomplish this, we first created a reduced design matrix *X*_r_, containing all movement regressors as well as times when spouts or handles were moving or visual stimuli were presented. *X*_r_ was ordered so that the motion energy and video columns were at the end. We then performed a QR decomposition of *X*_r_^60^. The QR decomposition of a matrix *A* is *A* = *QR*, where *Q* is an orthonormal matrix and *R* is upper triangular. Columns 1 to *j* of *Q* therefore span the same space as columns 1 to *j* of *A* for all *j*, but all the columns are orthogonal to one another. Finally, we replaced the motion and video columns of the full design matrix *X* with the corresponding columns of *Q*. This allowed the model to improve the fit to the data using any unique contributions of the motion and video regressors, while ensuring that the weights given to other regressors were not altered. Note that the QR decomposition is limited to the original space spanned by *X*, and therefore cannot enhance the explanatory power of the video regressors - it can only reduce it.

Subsequently, we used the same approach to orthogonalize all uninstructed movement regressors against instructed movements. This was done to ensure that all information that could be explained by instructed movements could not be accounted for by correlated, uninstructed movement regressors. As described above, this can only reduce the explanatory power of the uninstructed movement regressors.

The following table provides an overview of all model variables and how they were generated:

| Variable name | Description | Regressor type | Category |
| --- | --- | --- | --- |
| Hindpaw | Piezo sensor below the animal | Analog + Event kernel | Movement |
| Handles (Left / Right) | Touch events from handle sensors | Event kernel | Movement |
| Licks (Left / Right) | Lick events from spout sensors | Event kernel | Movement |
| Pupil | Pupil diameter, extracted from face camera | Analog + Event kernel | Movement |
| Nose | Nose movements, extracted from face camera | Analog + Event kernel | Movement |
| Whisking | Whisker movements, extracted from face camera | Analog + Event kernel | Movement |
| Body | Average motion energy across all body camera pixels | Analog + Event kernel | Movement |
| Video | Video dimensions from both cameras (SVD) | Analog | Movement |
| Video ME | Video dimensions from motion energy in both cameras (SVD) | Analog | Movement |
| Time | All trials | Event kernel | Task |
| Choice | All leftward choice trials | Event kernel | Task |
| Previous choice | Every trial after a leftward choice trial | Event kernel | Task |
| Previous modality | Every trial after a visual trial | Event kernel | Task |
| Previous success | Every trial after a successful trial | Event kernel | Task |
| Success | All successful trials | Event kernel | Task |
| Water given | All frames after a reward was given | Event kernel | Task |
| Audio (Left / Right) | All frames after a left- or rightward auditory stimulus | Event kernel | Task |
| Vision (Left / Right) | All frames after a left- or rightward visual stimulus | Event kernel | Task |

When a design matrix has columns that are close to linearly dependent (multicollinear), model fits are not reliable. To test for this, we devised a novel method we call “cumulative subspace angles.” The idea is that for each column of the design matrix, we wish to know how far it lies from the space spanned by the previous columns (note that pairwise angles do not suffice to determine multicollinearity). Our method works as follows: (1) the columns of the matrix *X* were normalized to unit magnitude, (2) a QR decomposition of *X* was performed, (3) the absolute value of the elements along the diagonal of *R* were examined. Each of these values is the absolute dot product of the original vector with the same vector orthogonalized relative to all previous vectors. The values range from zero to one, where zero indicates complete degeneracy and one indicates no multicollinearity at all. Over all experiments, the most collinear regressor received a 0.26, indicating that it was 15° from the space of all other regressors. The average value was 0.84, corresponding to a mean angle of 57°.

To avoid overfitting, the model was fit using ridge regression. The regularization penalty was estimated separately for each column of the widefield data using marginal maximum likelihood estimation^61^ with minor modifications that reduced numerical instability for large regularization parameters.

### Variance analysis

Explained variance (cvR^2^) was obtained using 10-fold cross-validation. To compute all explained variance by individual model variables, we created reduced models where all regressors that did not correspond to a given variable were shuffled in time. The explained variance by each reduced model revealed the maximum potential predictive power of the corresponding model variable.

To assess unique explained variance by individual variables, we created reduced models for each variable where only the corresponding regressor set was shuffled in time. The difference in explained variance between the full and the reduced model yielded the unique contribution ΔR^2^ of that model variable. The same approach was used to compute unique contributions for groups of variables, i.e., ‘instructed movements’, ‘uninstructed movements’ or ‘task’. Here, all variables that corresponded to a given group were shuffled at once.

To compute the ‘task-aligned’ or ‘task-independent’ explained variance for each movement variable, we created a reduced ‘task-only’ model where all movement variables were shuffled in time. This task-only model was then compared to other reduced models where all movement variables but one were shuffled. The difference between the task-only model and this model yielded the task-independent contribution of that movement variable. The task-aligned contribution was computed as the difference between the total variance explained by a given variable and its task-independent contribution.

### PETH partitioning

Reconstructed trial averages (Figs. 5 and 6) were produced by fitting the full model and averaging the reconstructed data over all trials. To split the model into the respective contributions of instructed movements, uninstructed movements and task variables, we reconstructed the data based on that variable group alone (using the weights from the full model, without re-fitting) and averaging over all trials. To evaluate the relative impact of each group variable on the perievent time histogram (PETH), we computed a modulation index (MI), defined as

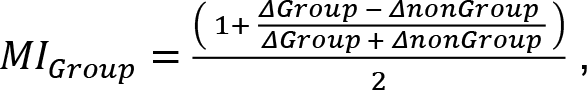

where ΔGroup and ΔnonGroup denotes the sum of the absolute deviation from zero when reconstructing the PETH either based on all variables of a given group (ΔGroup) or all other variables (ΔnonGroup). The MI ranges from 0 (variable group has no impact on PETH) to 1 (PETH is fully explained by variable group). Intermediate values denote a mixed contribution to the PETH from different group variables (Extended Data Fig. 9).

### Model-based video reconstruction

To better understand how the video related to the neural data, we analyzed the portion of the β-weight matrix that corresponded to the video regressors. This portion of the matrix was projected back up into the original video space. The result was of size *p* x *d*, where *p* is the number of video pixels (153,600) and *d* is the number of dimensions of the widefield data (200). We performed PCA on this matrix, reducing the number of rows. The top few ‘scores’ (projections onto the principal components) are low-dimensional representations of the widefield maps that were most strongly influenced by the video. To choose the dimensionality, we used the number of dimensions required to account for >90% of the variance (Fig. 3F). To obtain the widefield maps showing how the video was related to neural activity (Fig. 3G), we projected the scores back into widefield data pixel space and sparsened them using the varimax rotation. To determine the influence of each video pixel on the widefield (Fig. 3H), we projected the low-dimensional β-weights into video pixel space, took the magnitude of the β-weights for each pixel, and multiplied by the original standard deviation for that pixel (to reverse the Z-scoring step of PCA).

### Aberrant cortical activity in Ai93 transgenic animals

Mice with both Emx-Cre and Ai93 transgenes can exhibit aberrant, epileptiform cortical activity patterns, especially when expressing GCaMP6 during development^62^. To avoid this issue, we raised most of the 11 mice in our widefield data set (6 mice) on a doxycycline-containing diet (DOX), preventing GCaMP6 expression until they were 6 weeks or older. This was also true for all Ai93 mice in our 2-photon data set. However, 5 mice were raised on standard diet, raising the concern that aberrant activity may have affected our widefield results.

To test for presence of epileptiform activity, we used the same comparison as Steinmetz et al.^62^ on the cortex-wide average. A peak in cortical activity was flagged as a potential interictal event if it had a width of 60-235 ms and a prominence of 0.03 or higher. These parameters flagged nearly all cases of apparent interictal events (Extended Data Fig. 10A) and identified four out of the 11 mice in the widefield data set to exhibit potential epileptiform activity (Extended Data Fig. 10B). None of the identified mice were raised on DOX.

To ensure that epileptiform activity would not bias our results, we removed flagged events and interpolated over the resulting gaps (in low-D) with Matlab’s built-in autoregressive modeling (fillgaps.m) and a 20-frame prediction window. The result did not show any perturbations around the former interictal events (Extended Data Fig. 10C). When comparing modeling results between DOX- and non-DOX-raised mice, predicted variance was highly similar in all cases (Extended Data Fig. 10D-G). This shows that our results were not due to epileptiform activity and indicated that it was safe to include all mice in the dataset with this data processing step.

This was further supported by our additional experiments with GCaMP6s-expressing animals. Consistent with previous observations^62^, we found no epileptiform activity in these mice. Nevertheless, our model predicted an even larger amount of variance than in Ai93 mice and produced highly-specific maps of unique contributions from single variables (Extended Data Fig. 4).

