## Supplementary material for "Single-trial neural dynamics are dominated by richly varied movements"

**Allen common coordinate framework v3**

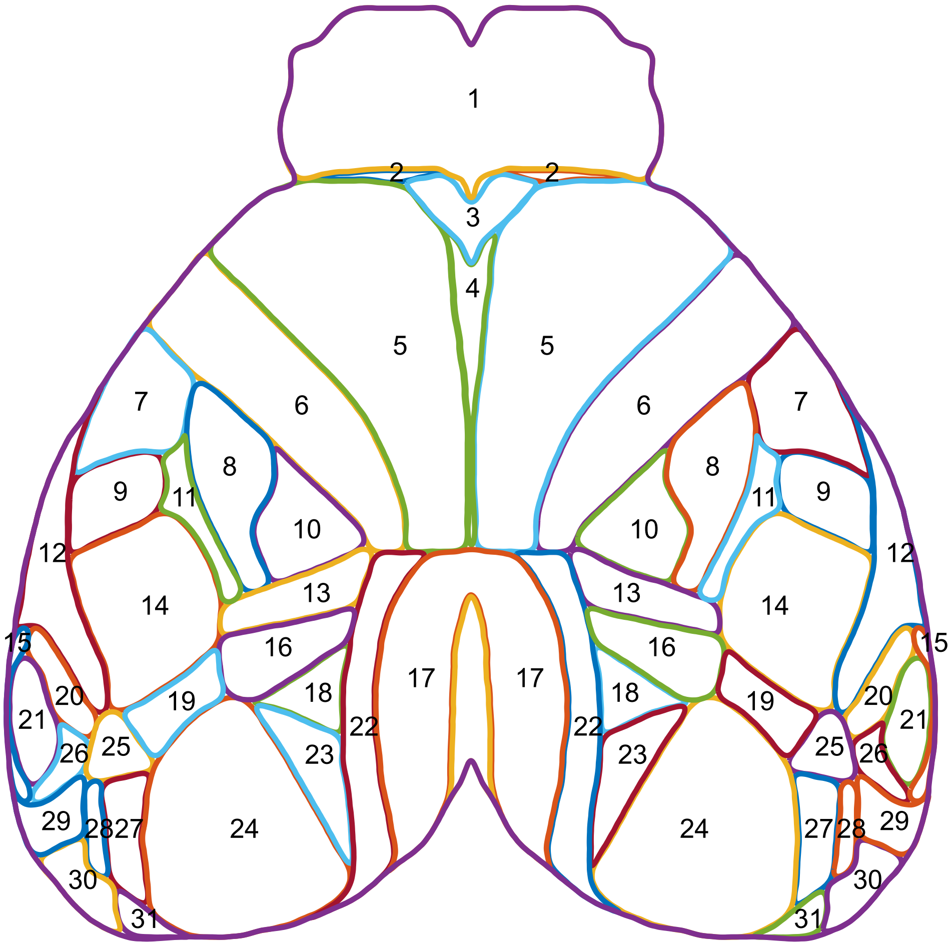


1 : Olfactory bulb (combined)
2 : Frontal pole, cerebral cortex
3 : Prelimbic area
4 : Anterior cingulate area, dorsal part
5 : Secondary motor area
6 : Primary motor area
7 : Primary somatosensory area, mouth
8 : Primary somatosensory area, upper limb
9 : Primary somatosensory area, nose
10 : Primary somatosensory area, lower limb
11 : Primary somatosensory area, unassigned
12 : Supplemental somatosensory area
13 : Primary somatosensory area, trunk
14 : Primary somatosensory area, barrel field
15 : Ventral auditory area
16 : Anterior visual area
17 : Retrosplenial area, dorsal part
18 : Anteromedial visual area
19 : Rostrolateral visual area
20 : Dorsal auditory area
21 : Primary auditory area
22 : Retrosplenial area, lateral agranular part
23 : Posteromedial visual area
24 : Primary visual area
25 : Anterolateral visual area
26 : Posterior auditory area
27 : Lateral visual area
28 : Laterointermediate area
29 : Temporal association areas
30 : Postrhinal area
31 : Posterolateral visual area

**Extended Data Figure 1. Overview over cortical areas.**
Shown are cortical areas based on the Allen common coordinate framework v.3.
The labels of the corresponding cortical areas are shown on the right.

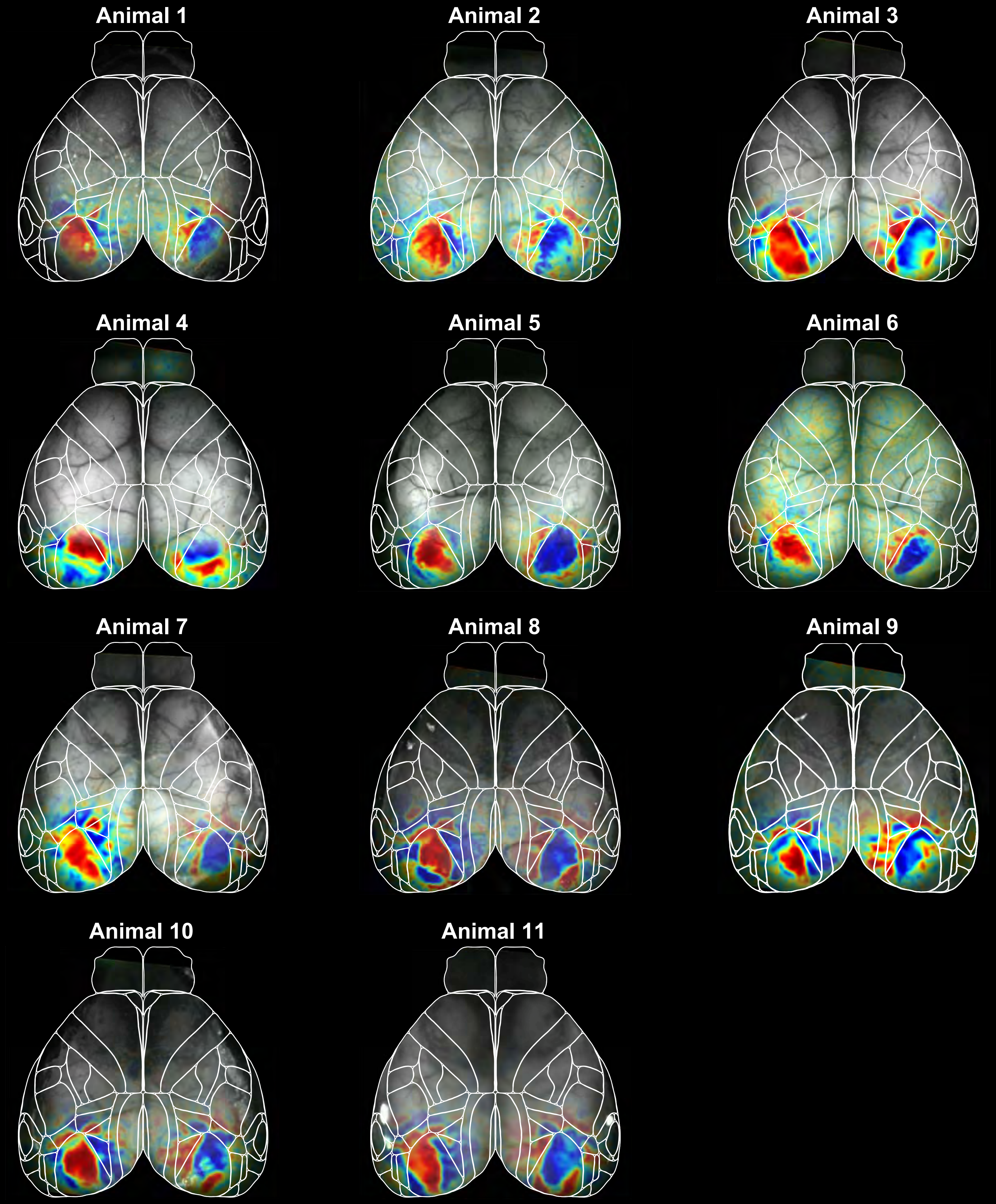


**Extended Data Figure 2. Visual sign maps for all mice.**
Shown are visual field sign maps for all trained animals, aligned to the Allen CCF. Mapped areas largely agreed with corresponding location of visual areas in the CCF.

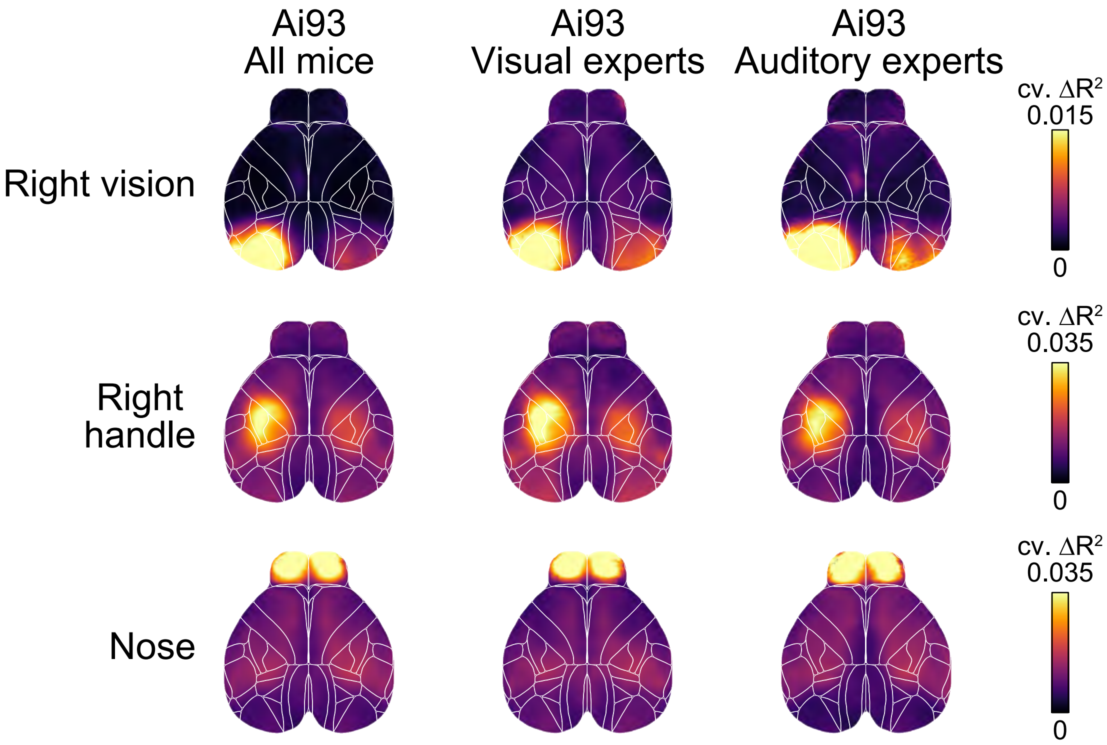


**Extended Data Figure 3. Unique explained variance for different expert groups.** Shown are cortical maps of unique model contributions for the right vision, right handle and nose variables. The left column shows the average over all recordings from 11 animals. All maps identified specific cortical areas that sensibly corresponded to their respective model variable. Maps were highly robust when averaging over all visual experts (6 mice, 12 recordings, middle column) or auditory experts (5 mice, 10 recordings, right column), respectively.

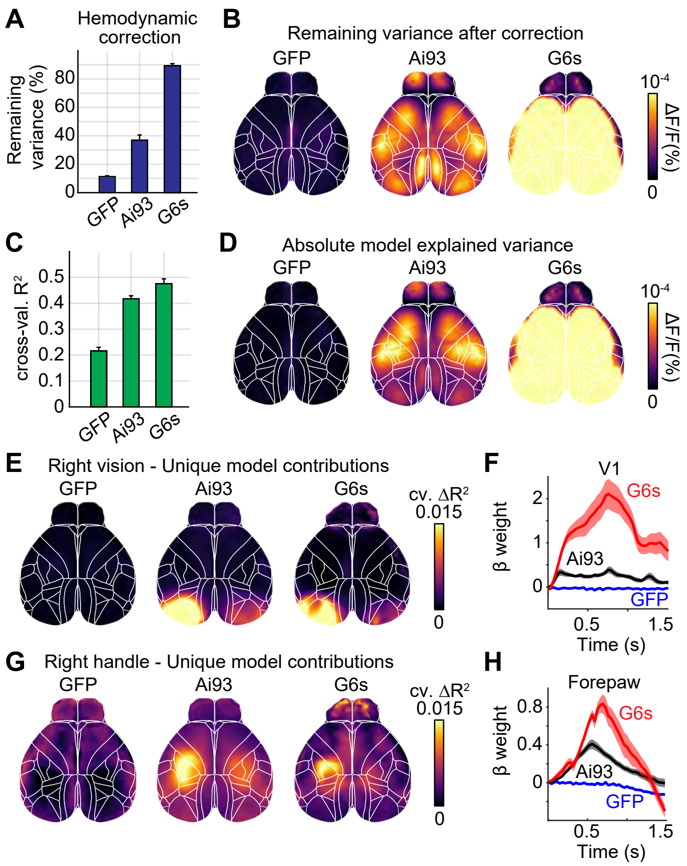


**Extended Data Figure 4. GFP and GCaMP6s controls.**
**(A)** Remaining variance in fluorescence after subtracting hemodynamic signals. Only ~10% of the variance remained in GFP controls (GFP), 38% in Ai93 mice (Ai93) and 89% in GCaMP6s expressing animals (G6s). This demonstrates that the hemodynamic correction accurately rejects most intrinsic activity while leaving calcium-related signals intact. **(B)** Absolute amount of remaining variance for individual pixels. Remaining variance in GFP controls was much lower as in GCaMP-expressing mice, across the dorsal cortex. **(C)** Cross-validated $R^2$ of the full linear model. Explained variance was lowest in GFP controls and highest in GCaMP6s animals, demonstrating that the widespread predictive power of the linear model is not explained by predicting hemodynamic signals. However, the model still accounted for ~21% of the variance in GFP animals, indicating that the hemodynamic correction is imperfect and the remaining fluorescence still contains a small but predictable component. **(D)** Absolute amount of predicted variance for individual pixels. Comparing absolute explained variance (instead of percentages in **A** and **C**) shows that, although a smaller percentage of fluorescence in GFP animals could be predicted, the absolute amount of predicted variance is extremely small compared to Ai93 mice. Absolute predicted variance was also much higher in GCamP6s-expressing animals, further demonstrating that the models success is due to accurate prediction of neural dynamics instead of intrinsic signals. **(E)** Unique model contribution maps for the right visual stimulus variable. No specific unique contribution was apparent in GFP mice but was clearly visible for Ai93 and GCaMP6s mice. Unique contributions were also well-localized to visual areas. **(F)** β-weights in V1. Visual responses are strongest in in GCaMP6s animals and absent in GFP controls. **G&H)** Same as **E&F)** for the right handle variable.

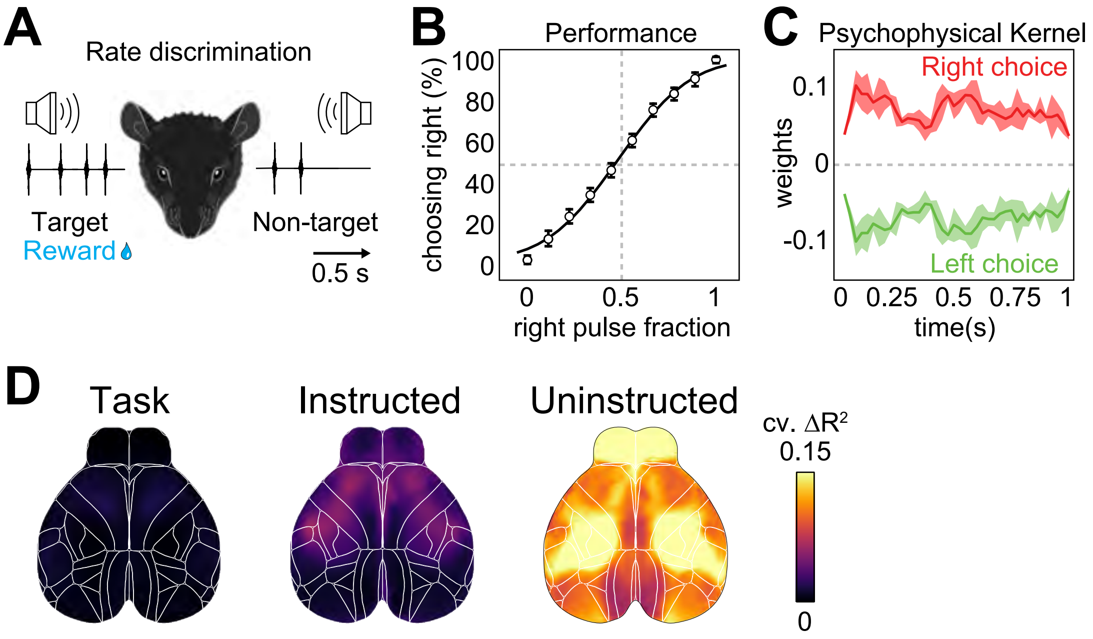


**Extended Data Figure 5. Auditory rate discrimination task**
**(A)** Schematic of the rate discrimination task. Mice are presented with auditory click sounds on both sides and identify the target side that contains more clicks to obtain a water reward. Stimulus sequences were 1-s long and clicks were randomly distributed. **(B)** Discrimination performance of an example animal. Right choice probability increases with the number of rightward pulses. Animals performed the task with high accuracy and were between 90-95% correct with the easiest stimuli. Shown are means ± 95% convidence intervals. **(C)** Psychophysical reverse correlation revealed time-points for which stimuli most strongly influence animal decisions. Positive weights predict rightward and negative weights leftward decisions. Weights were non-zero for the entire stimulus duration, demonstrating that animals integrated sensory evidence over time to perform the task. Shown are means ± s.e.m. for 4 mice. **(D)** Maps of unique model contribution for different variable groups, similar to Fig. 4E. As in the main results, unique contributions from uninstructed movements were highest across dorsal cortex. Shown are averages over 40 recordings from 4 mice.

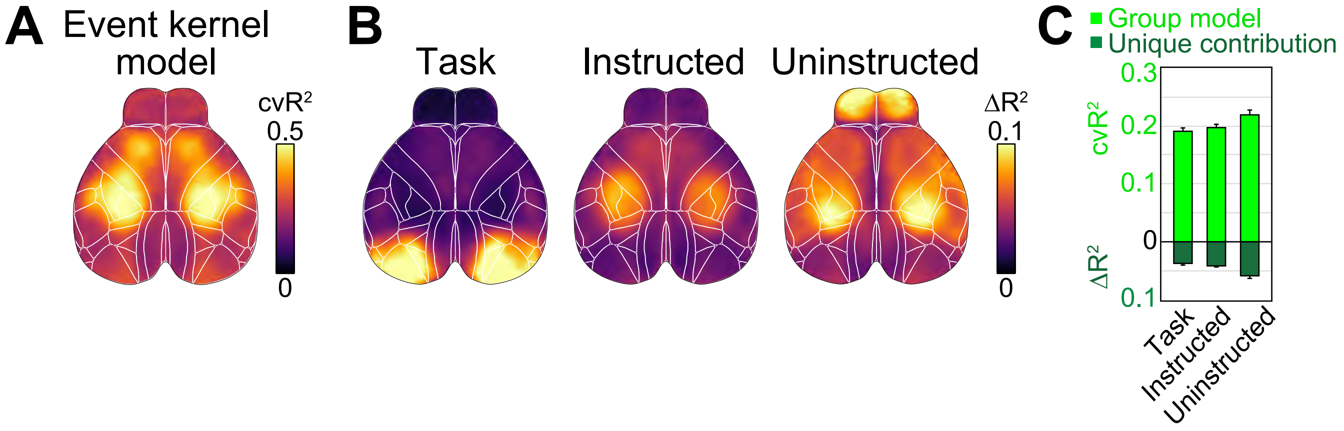


**Extended Data Figure 6. Model performance without analog predictors**

**(A)** Cross-validated explained variance for a reduced model without any analog regressors. Model's performance was lower than the full model in Fig. 3A but still predicted a large amount of variance. Averaged across cortex, the event kernel-only model predicted 30.8±0.2% (mean±s.e.m., n=22 sessions) of all variance. **(B)** Unique model contribution map for each variable group. **(C)** Explained variance for variable groups, averaged across cortical maps. Shown is either $cvR^2$ (light green) or $\Delta R^2$ (dark green). Bars represent mean ± s.e.m over 22 sessions. Even after removing all analog predictors, uninstructed movement contained the highest unique contributions across cortex. This demonstrates that their importance for predicting cortical activity is not just explained by including analog predictors such as the video variables.

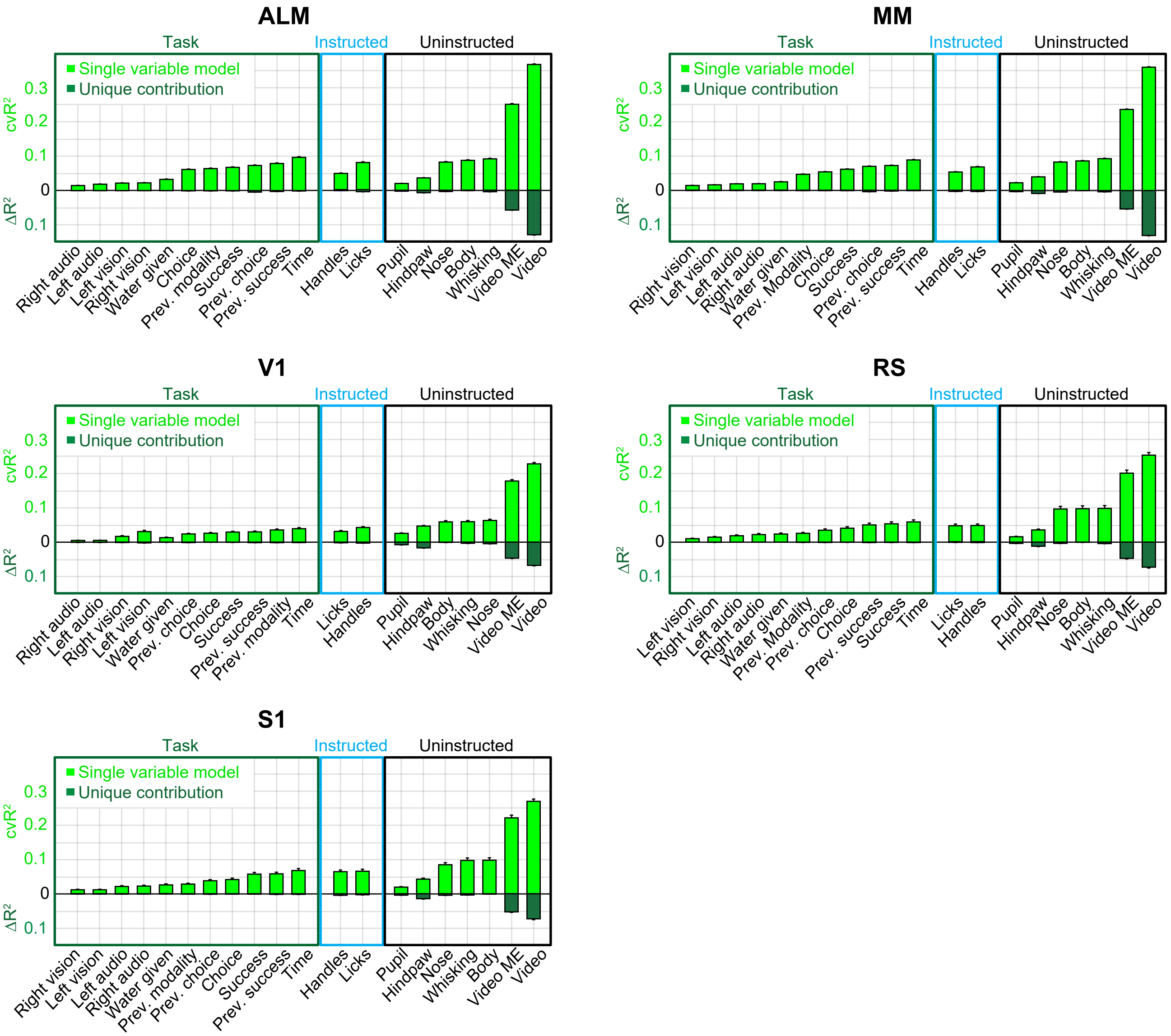


**Extended Data Figure 7. Predicted single-neuron variance for individual model variables**

Shown is either all explained variance ($cvR^2$, light green) or unique model contributions ($\Delta R^2$, dark green) for individual model variables. Values are averaged over all neurons in each recorded area. Bars represent mean ± s.e.m. Prev.: previous. n = 4571 neurons in ALM, 6364 neurons in MM, 594 neurons in V1, 206 neurons in RS and 252 neurons in S1.

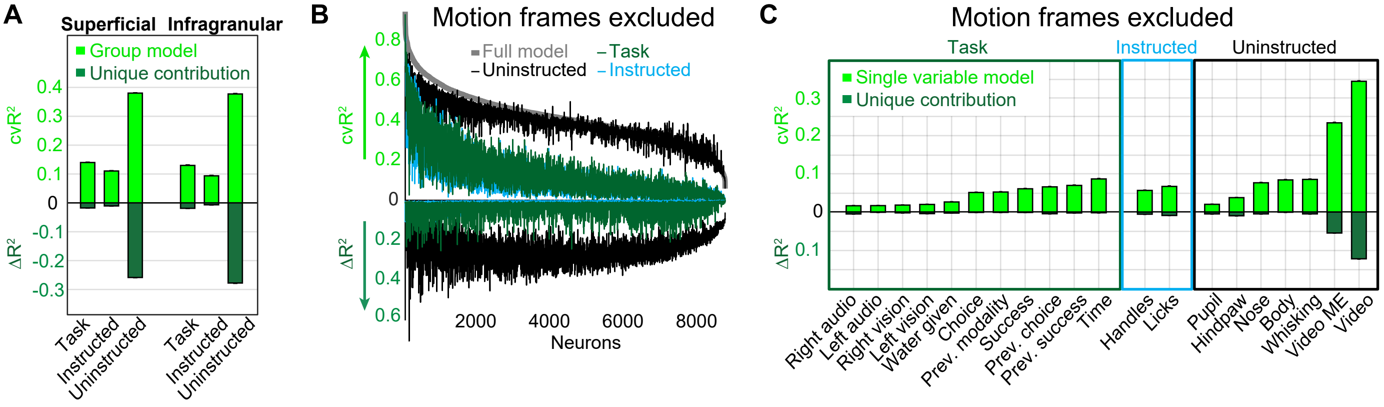


**Extended Data Figure 8. Depth comparison and motion control for 2-photon data.**
**(A)** Explained variance for groups of model variables at different cortical depths. Shown is either all explained variance (cvR$^2$, light green) or unique model contributions ($\Delta R^2$, dark green). Bars represent mean±s.e.m over 6923 neurons in superficial (left) and 6847 neurons in infragranular recordings (right). Superficial recordings were made from 150-350 µm, infragranular recordings between 350-450 µm. All infragranular recordings were performed in areas ALM (2655 neurons) or MM (3907 neurons). **(B)** Explained variance of variable groups for individual neurons, sorted by full-model performance (light gray trace). Conventions as in Fig. 6E but excluding imaging frames that were translated more than 2 pixels in either X- or Y-direction, relative to a reference image. As in Fig. 6E, uninstructed movements were most important to predict single-cell variance, demonstrating that this is not due to motion of the imaging plane with animal movement. **(C)** Explained variance for individual model variables averaged over all neurons after excluding translated imaging frames as described above. Shown is either all explained variance (light green) or unique model contributions (dark green). Bars represent mean±s.e.m over 8787 neurons.

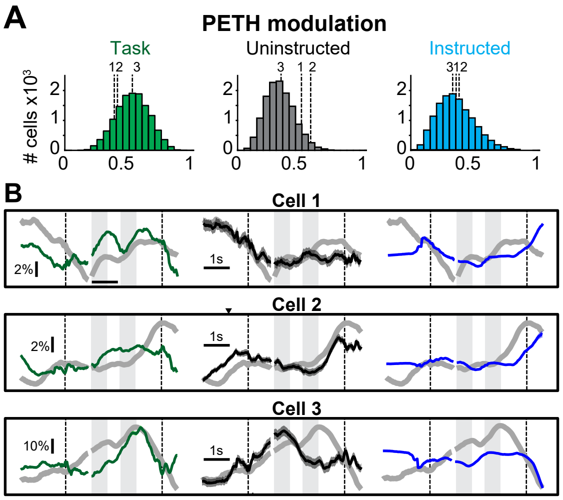


**Extended Data Figure 9. Revealing hidden task-dynamics with PETH partitioning**
**(A)** Histogram of modulation indices for either the task (left), uninstructed (middle) or instructed (right) movement groups. Dashed lines indicate the respective indices for the three example cells in **B**. In contrast to Figure 6H, all cells were best explained by a combination of all three model groups. **(B)** Three example cells with distinct task-related dynamics (left column, green) that were uncovered by accounting for uninstructed (middle column, black) and instructed movements (right column, blue). In all cases, task-related dynamics were clearly distinct from the uncorrected PETHs (gray traces) and revealed different response features, especially during the stimulus and delay period.

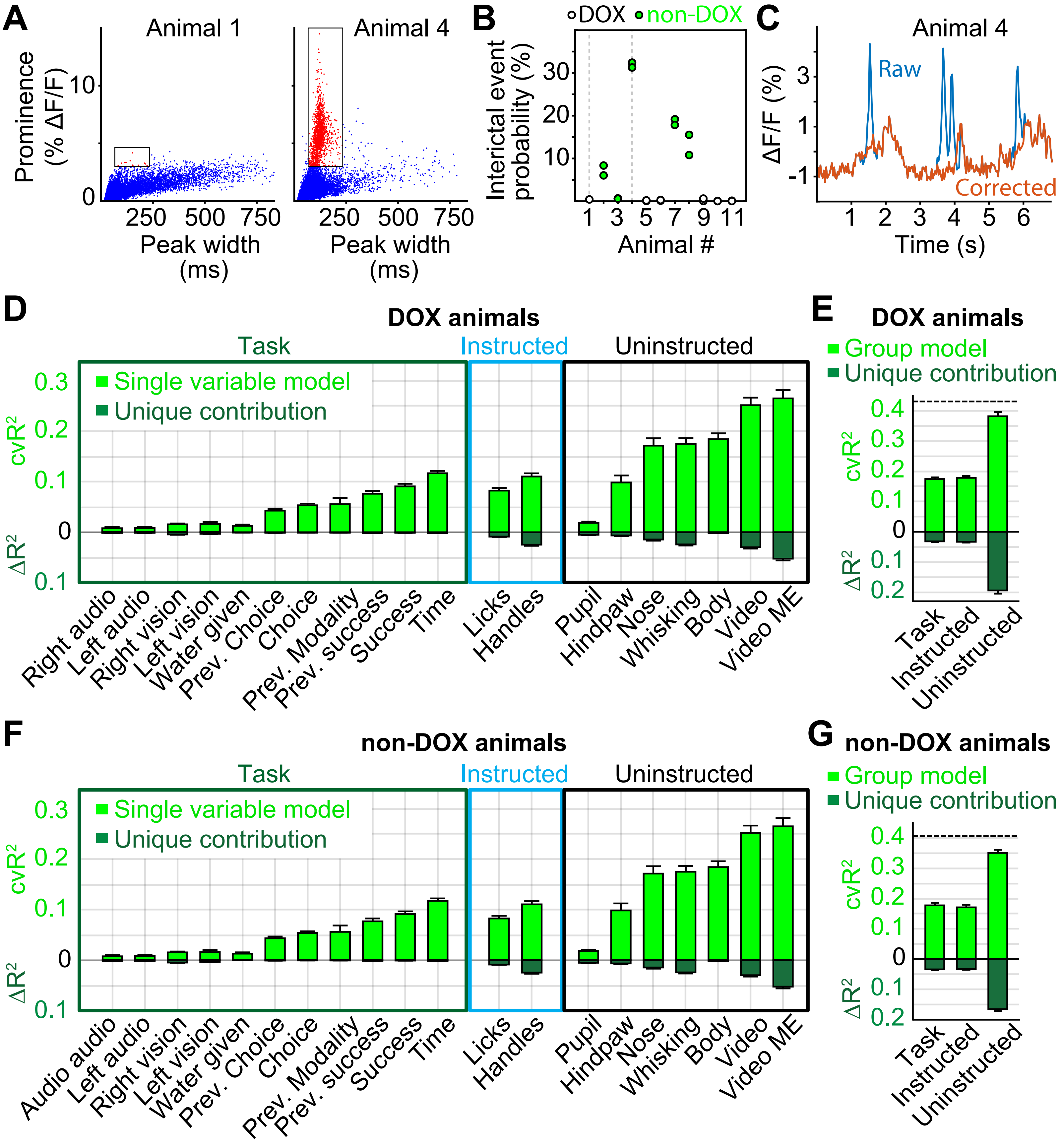


**Extended Data Figure 10. Controlling for interictal events.**
**(A)** Scatter plots show distribution of peaks in cortical activity, averaged over cortex. Left: Example animal, raised on a DOX-diet. Peaks were of variable length and remained at prominence below 5%. Right: Example animal, raised on a standard diet. Clearly visible are peaks of short latency and high promince (red dots). **(B)** Interictal event probability for all mice. Circles show individual sessions (two per animal). Four out of five mice that were raised on standard (non-DOX) diet show potential interictal activity. **(C)** Example trace for removal of interictal activity using autoregressive interpolation. **(D-E)** Modeling results for all DOX-raised animals. Similar to Figure 4 C,D. **(F-G)** Modeling results for all non-DOX-raised animals, showing potential interictal activity. Modeling results between DOX-raised and non-DOX-raised mice were highly similar, demonstrating that our results are not due to potential interictal activity in some of the mice.
